# A Spectral Image Scanning Microscope for Multi-Color High-Resolution Imaging

**DOI:** 10.1101/2025.05.14.653993

**Authors:** Lanna Bram, Ofir Tal-Friedman, Neta Fibeesh, Yohai Bar-Sinai, Eli Flaxer, Yael Roichman, Uri Ashery, Yuval Ebenstein, Jonathan Jeffet

## Abstract

Fluorescence confocal microscopy is a fundamental and widely used tool for biological, medical, and chemical research. It enables color-coded visualization of sample components, enhancing our understanding of life’s building blocks and their interactions. The recent realization of image scanning microscopy (ISM) using confocal spinning disks (CSD) enhanced the spatial resolution of fluorescence microscopy to twice the diffraction limit with minimal sample perturbation and fast acquisition rates. However, capturing multi-color images using ISM is still time-consuming and introduces temporal artifacts as different colors are not acquired simultaneously. Here, we present a spectral CSD-ISM system designed for concurrent high-resolution and simultaneous multi-color acquisition. By integrating a custom linear Amici prism into the CSD-ISM optical detection path, we achieve multi-color, super-resolution images at a fraction of the acquisition time and with a flexible color palette selection. A digital signal processor (DSP) is employed together with accompanying software as a cost-effective alternative to Field-Programmable Gate Arrays (FPGAs) used in previous studies. We provide an accompanying GPU-compatible, python-based image processing pipeline to decompose spectral signatures into multi-color channel-based images, preserving the optical resolution. System characterization using three color fluorescent beads demonstrated 1.74-fold resolution improvement over the diffraction limit and accurate color classification with three times faster acquisition compared to standard CSD-ISM. Application to neuron cells expressing a Parkinson’s disease-associated mutation, showcased improved resolution and contrast of four distinctly labeled cellular components. This spectral CSD-ISM system provides a valuable tool for biological imaging, enabling the simultaneous acquisition of high-resolution spatial information and multi-color spectral data.

## 1. INTRODUCTION

Science relies on fluorescence microscopy to visualize life at the smallest dimensions, enabling detailed visualization of cellular structures and dynamics. However, resolving intricate biological processes, especially within cells, necessitates observing the dynamic interactions between numerous molecules, often much smaller than the resolution limit (∼250 nm) of conventional optical microscopy. A common approach to enhance resolution is structured illumination [1,2], notably through Image Scanning Microscopy (ISM) [3]. In ISM, a focused laser beam scans the sample, akin to confocal microscopy [4,5]. However, instead of a point detector, a sensitive multi-pixel photodiode array or camera captures the emitted fluorescence, with its pixels functioning as an array of nearly zero-sized pinholes. This allows each pixel to simultaneously act as both a confocal pinhole and a detector, collecting the spatial distribution of emitted light around each illuminated point. The key innovation in ISM lies in the “pixel reassignment” process [6], where detected light is reassigned to its correct spatial position, effectively narrowing the point spread function (PSF) and improving resolution. This reassignment process with sufficient data sampling allows reconstructing a high-resolution image, achieving a two-fold improvement in lateral resolution compared to conventional confocal microscopy.

However, one significant drawback of the standard confocal ISM is its time-consuming nature. To address this, Schulz et al. [7] implemented ISM in combination with confocal spinning disk (CSD) [8,9], increasing image acquisition speed by nearly 100 times. Rather than a single pinhole, in CSD, two rotating disks, with thousands of pinholes and matching microlenses, spin at high speeds generating thousands of confocal excitation volumes that scan through the sample. Integrating over the entire disk rotation generates homogeneous illumination, with high contrast and Z-sectioning capabilities. In CSD-ISM, instead of integrating over the entire disk rotation, stroboscopic excitation is used at predefined disk rotation angles. This generates synchronized patterned illumination, which can be used with the ISM reassignment algorithm to double the resolution at a fraction of the acquisition time [10]. Consequently, the technique has become widely adopted in medical and biological research with commercial implementations of the technique available. Microscope modules such as SoRa (CSU-W1 SoRa Confocal Scanner Unit, Yokagawa) [11–14] and IXplore SpinSR (IXplore IX83 SpinSR, Olympus) [15–17] offer integral doubling of lateral resolution as their standard confocal module.

While high resolution is crucial, color information is equally important in fluorescence microscopy, providing essential contrast between different entities within the sample. Conventionally, multicolor imaging is established by sequential acquisition with different filters. In CSD-ISM, this leads to a significant extension of acquisition time, as hundreds of frames are needed in each channel to reconstruct the super-resolved image. Furthermore, this results in spatio-temporal chromatic artifacts, as different colors are recorded at different times [18]. Alternatively, split-view setups [19] allow for registering several colors simultaneously, albeit at the cost of a reduced field of view size and the requirement for cross-channel registration. More recently, spectral imaging was introduced [20–24], enabling the recording of color information unrestricted by discrete color channels. Spectral information is recorded by dispersing the detected light onto a different camera or a part of the camera. This, however, restricts throughput due to the large spectral dispersion induced, smearing each point spread function over a large pixel area, resulting in a relatively small effective FOV [25,26] (for further examples see Document S1. Supporting materials and methods, Table S1 in [27]). Recently, compact PSF engineering for spectral determination [27–32] has emerged. Methods such as continuously controlled spectral resolution (CoCoS) allow continuous tuning of the spectral dispersion for optimized performance [27]. These methods distribute color information locally, registering the information simultaneously for all colors with higher throughputs and higher fluorophore densities [27–29].

Here, we combine CSD-ISM with compact spectral registration. CSD-ISM, capturing a sparse array of confocal spots in each frame (Fig. 1b, left panel), offers a unique advantage for integrating spectral registration. By using a prism to disperse the spectral information contained in these spots, we encode all colors in a single CSD-ISM acquisition, better exploiting the full FOV of the camera (Fig 1b, right panel). This novel approach provides a powerful tool for high-resolution, multi-color imaging, enabling the simultaneous capture of spatial and spectral information by using a standard CSD microscope with minimal modifications.

**Figure 1.**
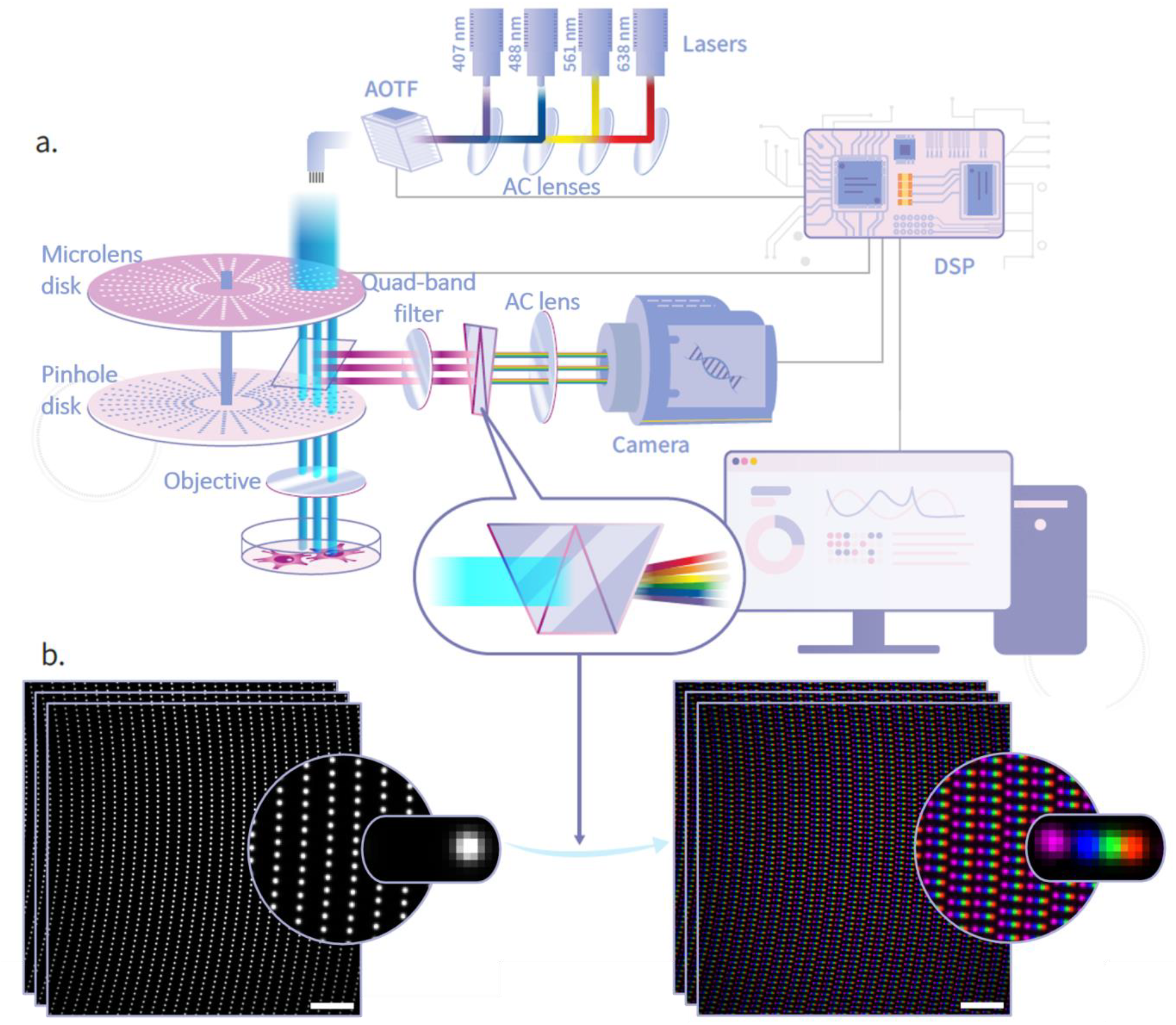
Schematic representation of the spectral Confocal Spinning Disk Image Scanning Microscopy (CSD-ISM) system. **(a)** *Optical setup*. Multiple laser lines are combined and controlled by an Acousto-Optic Tunable Filter (AOTF) to selectively excite fluorescence. The excitation light is directed through a spinning disk system comprising microlens and pinhole disks. The emitted fluorescence signal is collected through a polychroic mirror and a notch emission filter and passed through a custom Amici prism, which disperses the fluorescence emission into its spectral components and onto the camera. For the stroboscopic illumination required for ISM implementation, a Digital Signal Processor (DSP) reads the disk’s spinning frequency and sends pulsed signals to the AOTF and camera, synchronizing the laser pulses and camera exposures to the spinning disk frequency. **(b)** *Example imaging output from the system*. Image acquisition consists of several frames, each capturing a scanned sample image at a specific rotation angle of the disk. Summing all frames reconstructs the full image. The left panel shows raw images scanned without a prism, showcasing the structured illumination pattern induced by the disks. The right panel shows the result of dispersing colors using the custom amici prism. Each color is translated along the horizontal axis, capturing its spectral information. For illustration purposes, a false-colored, overlayed, multi-sample, and multi-acquisition (Alexa 405 in purple, Alexa 488 in blue, Alexa 568 in green and Alexa 647 in red) image was acquired using the custom prism (actual multi-fluorophore samples would have been recorded in a single grayscale manner). Scale bars are 10 μm.

## 2. PRINCIPLES

### a. Optical Setup

The optical setup was designed and constructed by combining spectral registration and ISM with a standard commercial CSD (figure 1a, for further details see ‘materials and methods’). To record high-resolution images, the stroboscopic illumination required for implementing ISM was controlled using a custom designed digital signal processor (DSP) [33,34]. The DSP synchronizes the spinning disks’ rotation frequency with the camera exposure, and the laser’s AOTF acquisition protocol was designed following the protocol outlined by Qin et al. [35], offering a cost-effective solution compared to the previous FPGA implementation (more than 70× price reduction [33]). A custom Amici prism was introduced at the output of the commercial CSD to register color simultaneously. Amici prisms, also known as direct vision prisms, disperse colors without deviating the optical axis [32,36], allowing convenient placement of the prism without requiring any additional optics. Unlike standard prisms which disperse shorter wavelengths more strongly, we especially designed the prism to disperse the visible spectrum linearly (supplementary section 1) in the vicinity of the CSD spots. The linear dispersion offered better exploitation of the area between illumination spots essential for maximizing color content. The integration of the prism results in the spectral dispersion of the fluorescence emitted from the sample, allowing us to encode each color into a unique spatial intensity distribution, which we termed the spectral PSF. Therefore, each confocal spot reports both on the fluorescence intensity and the underlying fluorescent color composition in its diffraction-limited volume (figure 2a).

**Figure 2.**
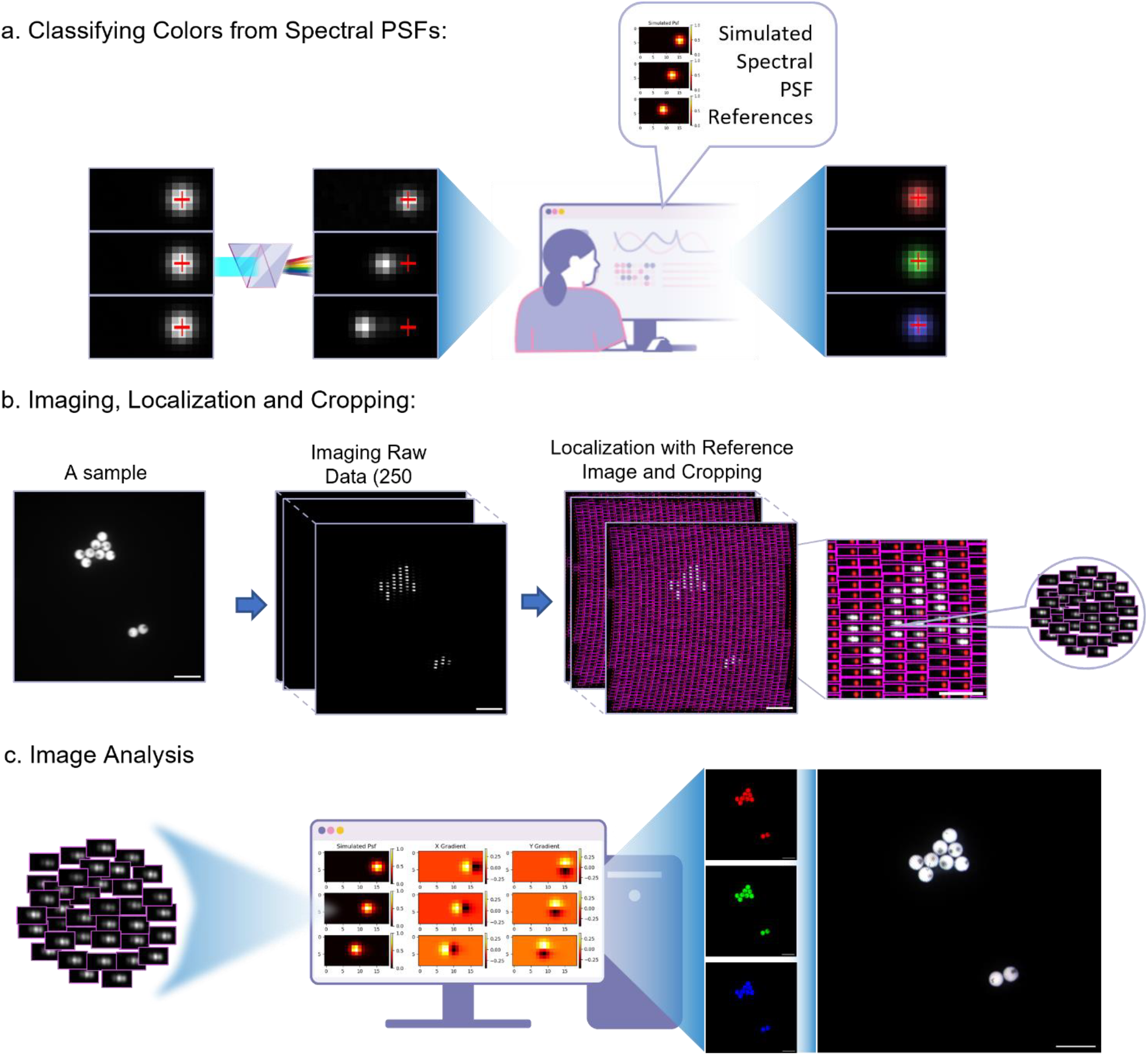
Workflow of CSD-ISM image processing, localization, and analysis. **(a)** *Schematic illustration of color classification*. Left: three simulated examples of confocal volumes containing three distinct colors without dispersion. Red crosses correspond with the confocal illumination reference localization. Center: a prism displaces each color with a distinct distance compared to the illumination reference, providing a unique spectral signature. Right: using linear regression with simulated spectral PSFs, the underlying color composition in each confocal volume is resolved and assigned to the non-dispersed location. **(b)** *Imaging, localization, and cropping process*. Left: Experimental image of three-colored 4 µm fluorescent beads without dispersion. Center: A stack of 250 frames imaged with spectral CSD-ISM is used to capture the full information content of the sample. Right: A reference stack of the illumination profile (in red) is used to crop spectral ROIs around each illumination volume across all frames. **(c)** *Image analysis and spectral decomposition*. Left: The spectral crops are decomposed into individual color channels using linear regression based on the simulated spectral PSFs and their gradients, and are spatially reassigned with the ISM algorithm. Right: The final output is a high-resolution multi-channel non-dispersed color image. Scale bars are 10 µm.

### b. Image Analysis

To accurately link fluorescence wavelengths to color readouts, establishing a conversion function between color and spectral PSFs is essential. This can be achieved through experimental imaging of homogeneous fluorescence samples to register their spectral responses or, more efficiently, via spectral PSF simulations. To create a conversion function between color and spectral PSF, we created a calibration curve using multiple homogeneously distributed fluorescent samples with emission peaks across the visible spectrum (figure 1 b as an example. for details, see methods and figure S2). For each sample, the spectral PSF’s center of mass was fitted with its corresponding emission spectrum’s center of mass after accounting for the spectral response of the camera and filters. This allowed us to establish a calibration model between spectral wavelength and pixel displacement, enabling the accurate simulation of any fluorescent spectral PSF according to its theoretical emission spectrum (see figure 2a and supplementary section 3, figure S3, S4). The resulting simulations not only validated the experimental data but also extended to unmeasured fluorophores, enhancing the versatility of the imaging system and serving as the linear basis for our downstream color classification model. Thus, this simple calibration, requiring no more than four samples across the spectrum, enabled us to extend our color palette and resolve fluorophores without the need for experimental registration of their spectral response.

After establishing the spectral response of our optical system, to capture multicolor high-resolution images, the frame-stacks of spectrally-encoded confocal spots need to be deciphered both spatially, using the ISM reassignment algorithm, and spectrally, converting the registered spectral signals into non-dispersed color-channel-assigned signal.

The ISM reassignment requires precise knowledge of the confocal spots’ illumination pattern with subpixel resolution at each frame [7,35]. To register their position, we used a reference sample of dense homogeneously distributed fluorescent far-red dye (Alexa Fluor 647 – AF647), localizing and registering all illumination spots’ subpixel positions in each frame (figures 2a and 2b) using our open-access GPU-enabled python image analysis pipeline (https://github.com/ebensteinLab). With a single calibration acquisition, we could easily implement the ISM algorithm on all samples acquired that day (see methods).

The color-from-spectral reassignment was carried out using the same reference. Essentially, each illumination confocal spot can excite a linear combination of fluorophores found within its confocal volume. Therefore, we used linear regression to estimate the linear combination of spectral PSFs imaged around each registered location in the reference. In practice, we crop a rectangular region of interest (ROI) from the sample’s data around each of the registered illumination localizations (figure2b). For each ROI we estimate its underlying fluorophore composition by comparing it to a linear combination of simulated spectral PSFs according to the dyes used to label the sample. Finally, we assign the extracted linear coefficients to a multi-channel non-dispersed PSF, creating a multi-channel color representation of the ROI positioned at the reassigned spatial location according to the ISM algorithm (figures 2a and d2c).

One key principle underlying the resolution enhancement of ISM is the parallax effect when imaging the same sample from different angles. Small paraxial spatial shifts induced by exciting and imaging the sample through a variety of microlens positions and viewing angles (see supplementary section 3, figure S3), allow with the correct signal reassignment to double the resolution. To approximate the parallax-induced spatial shifts during our color assignment, we extracted the first term coefficients of the two-dimensional Taylor expansion by adding the spectral PSFs’ gradients to the linear regression model (figure 2c center, detailed explanation in supplementary section 3). This allowed us to maintain the spatial resolution during the color reassignment with only a slight deterioration (figure 2c, supplementary section 3 and figure S5). A beneficial feature of the linear regression analysis used for color assignment is that it is calculated by a closed formula without the need for optimization of the coefficients. This allows calculating the color compositions of millions of cropped ROIs in a fraction of a second on the CPU, keeping the computation time on par with the imaging acquisition time. Finally, after each ROI was re-assigned both spatially and spectrally, the individual ROIs are summed and reconstructed to create a multi-channel composite image of the sample (figure 2c, right panel). Finally, for resolution enhancement, ISM reconstruction and further deconvolution were applied. A flowchart scheme of the full image analysis pipeline is presented in figure S6.

## 3. RESULTS

### a. Spectral CSD-ISM Characterization and Validation

To benchmark the color and spatial resolution of our spectral CSD-ISM system, we imaged a mixture of three samples of single-colored, sub-diffraction-limit, fluorescent beads (170 nm diameter, emission maxima at 515 nm, 560 nm and 660 nm for the blue, green, and red beads respectively) (see ‘Methods’ section for details). The results presented in figure 3 show good assignment of colors, and a clear improvement in the resolution of the ISM and the Lucy-Richardson [37,38] deconvolved ISM images (see ‘Methods’ for a description of the deconvolution process). To quantify the improvement in resolution, we measured the full width at half maximum (FWHM) of the red beads. These beads, unlike the other colors, presented almost no apparent dispersion in our system (see supplementary section 3, figure S4), thus allowing a direct comparison between images pre- and post-color and ISM reassignment. Our calculation (supplementary section 5, figure S8) shows a 1.74± 0.24-fold resolution enhancement of the deconvolved color ISM image compared to the sum of raw spectral images. As shown in figure 3c, the color assignment slightly deteriorates the resolution. This is because of the limited accounting for parallax effects with a single term of the Taylor approximation. Introducing a second Taylor term (second derivatives) to the linear least squares regression did not improve the results (see supplementary section 3, figure S5). Nonetheless, the ISM reconstruction and deconvolution significantly improved spatial resolution, overcoming this initial loss. Consequently, a single acquisition of 500 frames provided a high-resolution three-color image, eliminating the need to make multiple multi-frame acquisitions for each color separately as in standard CSD-ISM.

**Figure 3.**
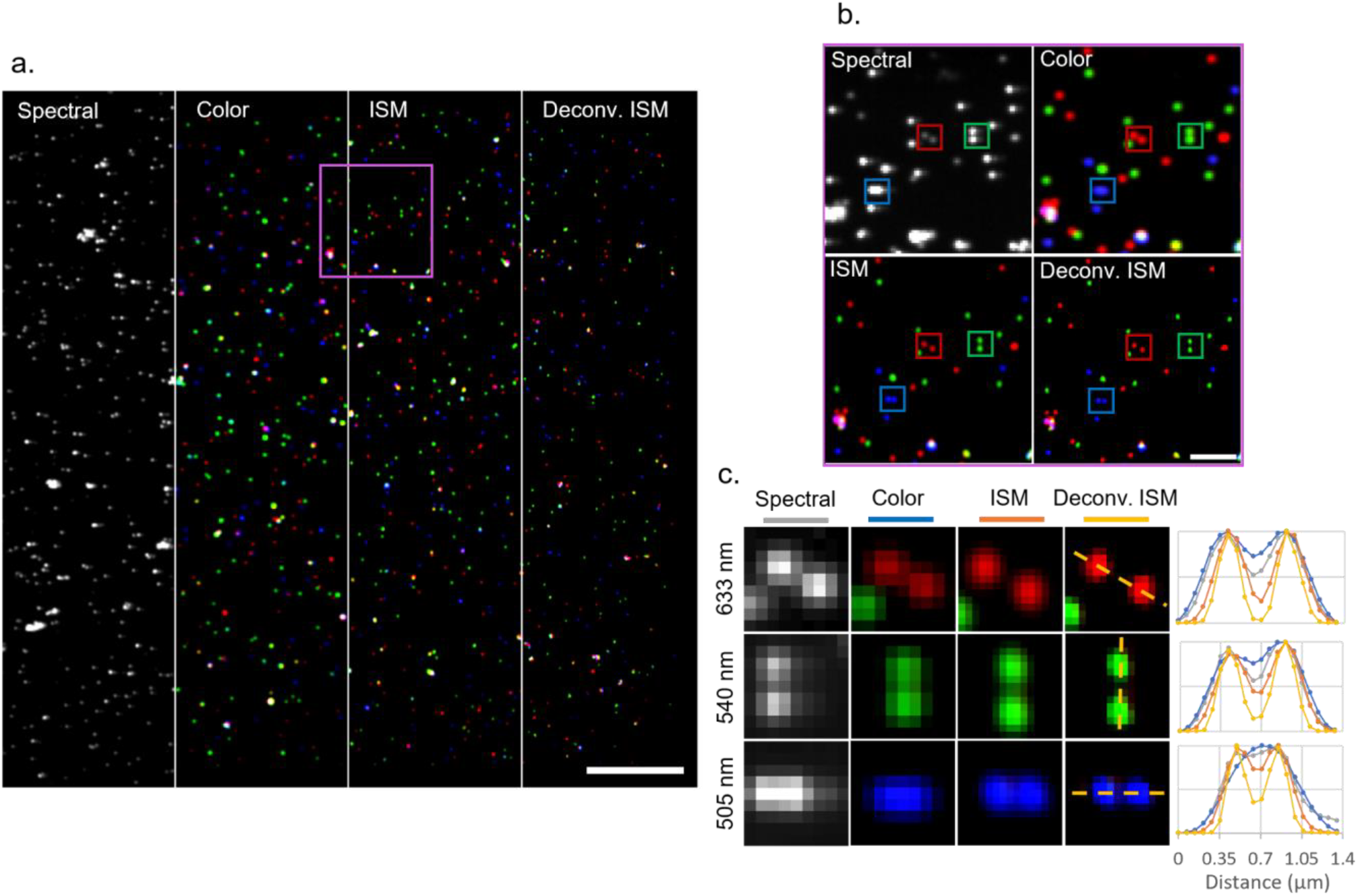
Spectral ISM Validation with Fluorescent Beads. **(a)** *Widefield image of 170 nm fluorescent beads with emission maxima at 515 nm (blue colored), 560 nm (green colored) and 660 nm (red colored)* at different stages of processing. The different sections of the image showing: an average of 500 spectral frames (spectral), color-assigned no-dispersion image (color), color ISM image (ISM), and the final deconvolved ISM image after applying Lucy-Richardson deconvolution (Deconv. ISM). Scale bar: 10 µm. **(b)** *Magnified view of the magenta rectangle highlighted in panel (a)*, demonstrating the resolution enhancement. Rectangles mark zoomed-in regions shown in panel c. Scale bar: 2 µm. **(c)** *Representative examples of pairs of beads of different colors* at each processing stage. At the right side, Normalized intensity profiles of the bead pairs along the dotted yellow lines.

### b. Biological Samples

To showcase our method in a biologically-relevant scenario, we imaged neuron cells expressing a mutant asynuclein (a-Syn) associated with familial Parkinson’s disease [39]. Neurons were transduced with an adeno-associated virus consisting of the mutation and a green fluorescent protein (GFP) label to investigate specific downstream consequences of pathological a-Syn on proteins of interest. Such proteins include the RNA binding protein G3BP and the synaptic protein synaptotagmin-1. G3BP marks stress granules which can range in sizes from 100 nm to several micrometers [40] and synaptotagmin-1 is a key component of synaptic vesicles which can be as small as 40nm [41]. Whilst diffraction-limited microscopy is limited to 200 nm resolution, improved visualization and deconvolution of these structures can facilitate enhanced understanding of the localization, interaction and aggregation of such structures. Apart from the endogenous GFP, immunofluorescence was used to fluorescently label the proteins G3BP and synaptotagmin-1 with CF568 and CF647 secondary antibodies, respectively. DAPI was used to stain DNA in the nuclei. Figure 4 presents an image from a neurite portion of a cell labeled with eGFP, and shows a clear enhancement in resolution and contrast. Some ISM reconstruction artefacts are visible in the deconvolved image. Due to the low excitation power in our system (see methods) these artefacts are more pronounced, and become more visible with reducing the number of acquired frames per ISM image (see figure S10). These were characterized both theoretically and experimentally in previous works by the Enderlein group [35,42].

**Figure 4.**
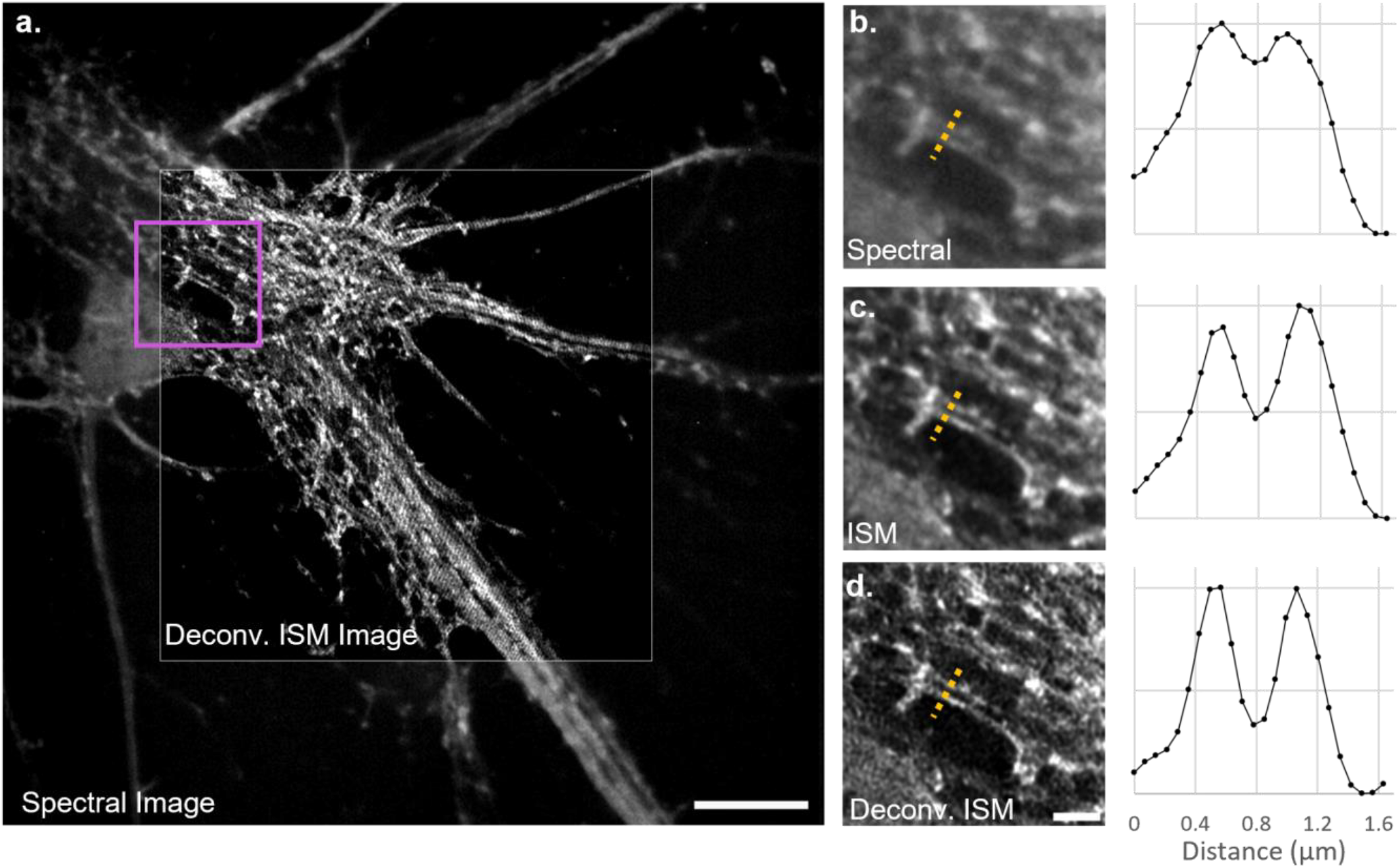
Spectral CSD-ISM Enhances Resolution in Neuron Imaging. **(a)** *Image of adeno-associated virus endogenous eGFP expression in a neurite portion of a neuronal cell*. An image composed of the sum of spectral frames is in the background, and a deconvolved single-color, non–dispersed, ISM image is in the center. The purple box indicates the region of interest in b-d. Scale bar, 10 µm. **(b-d)** *Resolution comparison in the selected region with corresponding normalized intensity profiles*: (b) Sum of spectral images (Raw image), (c) ISM image, and (d) deconvolved ISM image. Yellow dotted lines mark the profile measurement locations. scale bar 1 µm.

Figure 5 presents simultaneous four-label imaging with spectral CSD-ISM imaging with a single spectral acquisition of 250 frames for the four-color ISM image. The improvement in resolution is evident, revealing finer cellular structures. The zoomed-in images reveal some artifacts caused by the low frame count and SNR.

**Figure 5.**
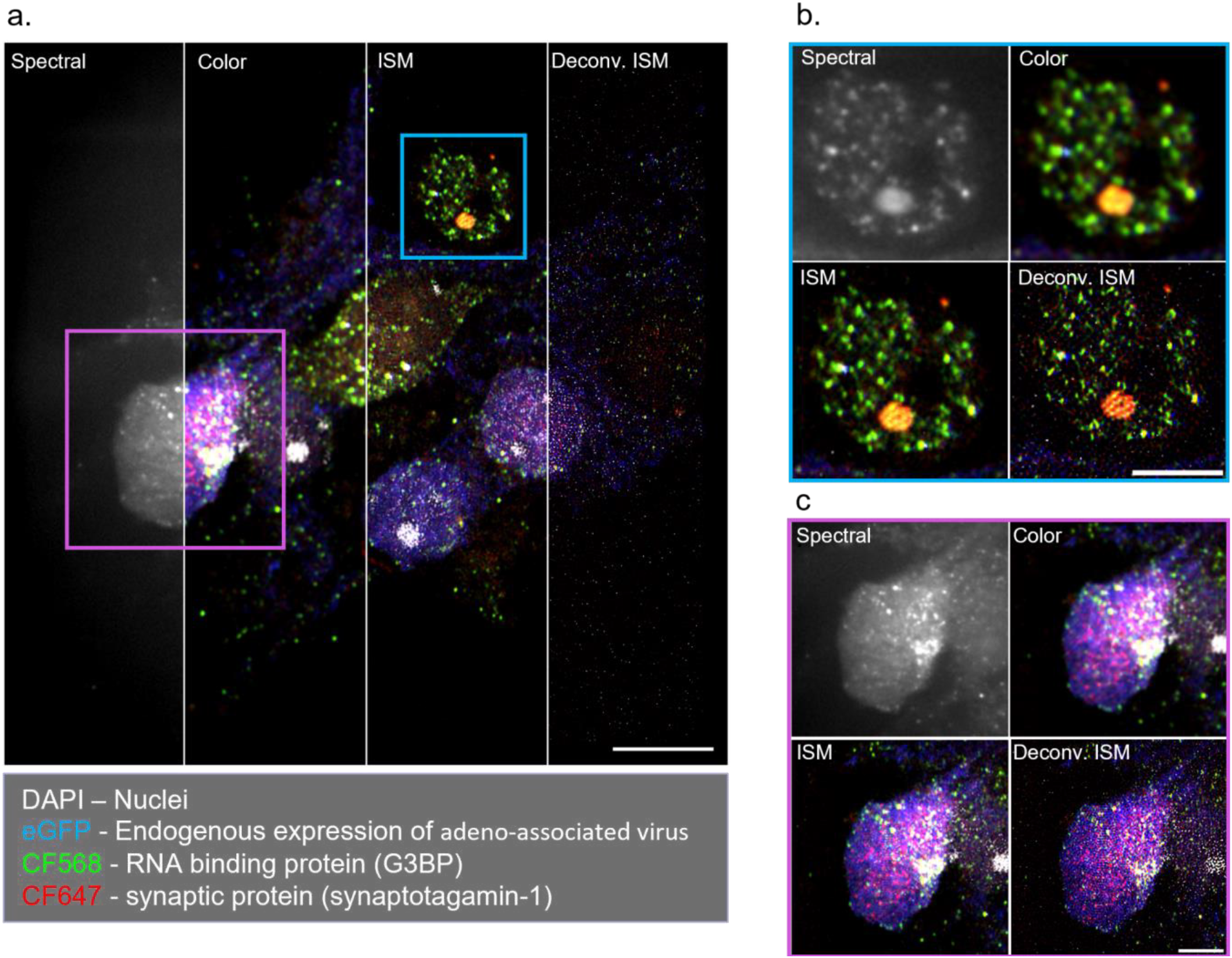
Spectral ISM Imaging of Biological Cells. **(a)** *Widefield images of biological cells at different processing stages*: sum of 250 spectral images (spectral), sum of color assigned image without ISM reassignment (color), color assigned with ISM pixel reassignment (ISM), and RL-deconvolved spectral ISM image (deconv. ISM). Color code: white - DAPI stained DNA in the nuclei; blue - endogenous expression of eGFP by adeno-associated virus; green - CF568 labeled G3BP RNA binding protein; red - CF647 labeled synaptotagmin-1 synaptic protein. Scale bar, 10 μm. **(b**,**c)** *Zoomed-in views of selected cellular regions*, as indicated in (a). Scale bar, 5 μm.

## 3. CONCLUSIONS AND DISCUSSION

In this study, we have successfully developed and validated a novel spectral CSD-ISM system that combines high-resolution imaging with spectral acquisition, enabling the simultaneous recording of spatial and color information in biological samples. By integrating a custom-designed linear Amici prism into a CSD-ISM setup, we achieved efficient spectral encoding and improved resolution, as demonstrated through imaging of fluorescent beads and neuron cells.

To convert the acquired spectral information into standard color-channel readout, we have established a comprehensive pipeline (see flowchart in figure S6) coded in python and openly available via github (https://github.com/ebensteinLab). The experimental protocol requires two calibration steps: first, a one-time experimental registration and calibration of the system’s spectral response, allowing us to computationally extend the spectral readout to any fluorescent marker with known emission spectrum; and second, an experimental illumination pattern registration, recorded once per day, to accurately calibrate the ISM algorithm. With these two calibrations, the full experimental spectral data stack (acquired in one-go) is segmented into ROIs, which subsequently undergo spectral-to-color reconstruction using a fast linear least squares regression. The reconstructed color-channel ROIs are then summed using the ISM pixel-reassignment algorithm creating a high-resolution color-channel image.

Our system characterization using multi-colored 4 μm fluorescent beads confirmed the accurate color representation and validated the image reconstruction process. Furthermore, imaging of 170 nm (sub-diffraction-limit) fluorescent beads demonstrated a 1.74-fold resolution improvement from the spectral image to the deconvolved spectral ISM image, highlighting the effectiveness of our approach in enhancing spatial resolution.

Our method offers a boost to the image acquisition speed since it enables simultaneous imaging of all channels, rather than capturing individual color-channels sequentially. However, since our linear regression model is limited in registering large parallax-induced lateral translations, we observed a slight decrease in lateral resolution. As discussed in supplementary section 3, further improvements to the resolution could be introduced by improving the regression analysis, thus accounting for larger parallax translations without interfering with color assignment. Nonetheless, our approach offers a good balance between performance and resolution, allowing us to analyze millions of spectral ROIs in a fraction of a second on the CPU with only a slight deterioration of resolution (figure S8). Maintaining a fast calculation is advantageous as it allows for a future implementation as an on-the-fly process, where the color decomposition is calculated alongside with the spectral image acquisition.

Another limitation of our current process is the requirement for non-overlapping spectral PSFs for accurate color decomposition; samples must be prepared with fluorophores who’s spectral PSFs are separated by at least the FWHM of the PSFs, to ensure correct color differentiation. Although this is a standard requirement in fluorescence microscopy to avoid bleed-through between channels, we have strong evidence [31,32] that future works using machine learning could exploit subtler nuances of the spectral PSF, thus overcoming this limitation. The application of spectral CSD-ISM to image neuron cells from a Parkinson’s disease study showcased the system’s ability to reveal finer cellular structures and improve image contrast. Although some artifacts, notoriously known in CSD-ISM imaging [42], were observed due to low signal-to-noise ratio (SNR) and frame rate limitations, these can be addressed by further optimizing imaging parameters and hardware alignment. To summarize, this integrated approach offers several advantages. First, it shortens and simplifies the process of acquiring multi-color images, eliminating the need for color alignment by computationally intensive post-processing. Second, it leverages the inherent super-resolution capabilities of CSD-ISM, allowing for the visualization of fine details within biological samples. Finally, it is readily implementable in any lab equipped with a CSD system, making high-resolution spectral imaging more accessible to a broader range of researchers. Overall, the spectral CSD-ISM system presented here offers a powerful tool for multi-color, high-resolution imaging of biological samples. Future work will focus on optimizing the system for improved SNR, reducing artifacts, and expanding its capabilities for 3D imaging and dynamic live-cell applications.

## 4. MATERIALS AND METHODS

### a. Experimental Setup

Initially, we reconstructed the CSD-ISM system much like the protocol outlined by Shun Qin et al. [35]. The essential components of our CSD-ISM include an Olympus IX71 Inverted Research Microscope implemented with a 100×, 1.40 NA objective lens (Olympus UPlanSApo 100x, Oil) that was controlled by a piezo (E-665 Piezo Amplifier / Servo Controller), Andor Precision Control Unit ER-PCUB-110, and Yokogawa Confocal Scanner Unit CSU-X1. As a light source, we used Andor Laser Combiner ALC-501A, with laser wavelength and power modulation via an AOTF that contained four lasers-407 nm (Coherent CUBE), 488 nm (Sapphire Coherent Laser), 561 nm (Cobolt Jive) and 638 nm (Cobolt 06-MLD). Throughout all measurements, the lasers were operated with a laser power at the sample plane of 0.02 mW, 0.45 mW, 0.22 mW and 0.5 mW, respectively. A multi-band filter (440/521/607/700 nm BrightLine® quad-band bandpass filter) was used in the detection path of the CSD. For dispersion, a customized linear Amici prism was used and integrated between the CSD and the camera. Final images were collected by an Andor iXon+ 897 emCCD Camera. For hardware synchronization, we used a digital signal processing (DSP)-based controller integrated with a Windows-based control software [33]. The DSP receives an input signal from the spinning disk control unit (CSU) and sends two digital output signals to the camera and the AOTF.

### b. DSP-based Controller

For hardware synchronization in our spectral CSD-ISM system, we implemented a digital signal controller (DSC). Based on the methodology in [33], the DSC was selected as a cost-effective and accessible alternative to FPGA solutions. The DSC synchronizes the CSD frequency with the laser pulse (managed by an AOTF) and the camera shutter. The DSC receives pulses from the CSU as a reference and then generates two digital waveforms: one for the camera shutter and the other for laser activation, with each rising edge of the input pulse initiating a new scan cycle. Furthermore, a Windows-based application was provided for simplified control of the system’s triggering hardware. This software manages eight digital parameters and four analog laser intensity values through a custom communication protocol. the parameters are:

- TC_ON: Spinning Cycle per Frame
- TC_OFF: Spinning Cycle for Camera Readout
- TL: Laser Pulse Width (µs)
- TR: Laser Pulse Rate (µs)
- TD: Frame Shift Time (µs)
- TA: Camera Activation Time (µs)
- LPN: Laser Pulses per SSC (Single Scan Cycle)
- N: Total Frames’ Number

The DSP calculates a dynamic delay time (TK) for each frame using the formula: TK = TA + K*TD, K is the frame index, ensuring accurate temporal coordination during image acquisition. Furthermore, it also monitors the rotating disk frequency, enforcing adherence to predefined parameter limits. An automated rate function measures the disk frequency, calculates TR and TD values, and automatically configures the hardware accordingly. To evaluate the effectiveness of CSD-ISM with this DSP, we benchmarked it with a sample of diffraction-limited fluorescent beads and confirmed a resolution enhancement of x2.11, in accordance with ref. [35] (see figure S7). For three-dimensional (3D) imaging, further settings need to be established, including the total number of layers for 3D imaging, specifying the interval between layers, and setting the initial position (figure S11). The application and its communication library were developed using National Instrument LabWindows/CVI, ensuring compatibility with ANSI C and the Win32 API.

### c. Prism Design

To extend the multiplex capabilities of our system we designed a custom compound double Amici prism. Amici prisms retain the optical axis for a central wavelength while light of shorter or longer wavelengths is dispersed in opposite directions. We optimized our prism design using Zemax [43] for distinguishing between five spectral channels by exploiting the available pixel area between adjacent CSD excitation spots (figure S1). To efficiently encode all spectral channels, we designed the prism’s dispersion to have a linear wavelength dependence, ensuring homogenous spectral resolution for the entire visible spectrum and therefore optimized multiplexing (additional information on the optimization and design is given (supplementary section 1). Based on this optimization, a linear Amici prism was manufactured (Shanghai Optics) and integrated into our setup with a custom built, light-tight holder at the output of the CSU-X1 (Yokogawa).

### d. Sample Preparation

#### i. Spectral Calibration

Homogeneous emission across the field of view is essential for acquiring reference images for calibration at different wavelengths. We used a variety of dyes each dissolved in DMSO to 10mM concentrations (Alexa Fluor (AF) dye 405-DBCO, AF488-DBCO, AF532-DBCO, AF546-DBCO, AF568-DBCO, AF647-DBCO, and Cy5.5-DBCO, Jena Bioscience, Jena, Germany), and diluted at 1:100 volume ratio with TE buffer (10 mM Tris and 1 mM EDTA (pH 8)). 8 µl of the resulting solution was placed between a prewashed microscope slide and a standard 22×22 mm^2^ coverslip in order to create a homogeneous dye solution film for each color.

#### ii. Image Analysis Pipeline

We used a commercially available test slide containing 4 µm multi-colored fluorescent beads, stained with blue (350/440 nm), green (505/515 nm), orange (540/560 nm), and deep red (633/660) dye coatings (FocalCheck fluorescence microscope test slide #1, slot A3, ThermoFisher).

#### iii. Spectral CSD-ISM Characterization and Validation

For validating the resolution and color reconstruction we used sub-diffraction limit, 170 nm fluorescent beads (green (505/515 nm), orange (540/560 nm), and deep red (633/660) PS-Speck microscope point source kit; Thermo Fisher). To prepare the slide, we mixed 2µl bead solution of each color with the kit’s mounting medium and sonicated the mixture for 10 minutes. Then, 3 µl of the final mixture was placed on a microscope slide, air-dried for a few minutes, and covered with a clean 22×22 mm^2^ coverslip.

#### iv. Biological Samples

For cell imaging we used primary mouse neuron cells expressing a mutant a-synuclein (a-Syn) associated with familial Parkinson’s disease via viral transduction. The neurons were prepared from D0-D1 mouse pups in accordance with the ethical guidelines of Tel Aviv University. Briefly, neurons cultured for 14 days and were fixated with 4% PFA, washed with 1X PBS, permeabilized with 0.1% Triton-X then blocked with bovine serum albumin (BSA) and normal goat serum (NGS). Next, neurons were incubated with primary antibodies G3BP (1:250 Abcam ab181150) and synaptotagmin-1(1:500 synaptic systems #105001) at 4 degrees Celsius overnight, followed by 1X PBS washes and then CF568 (1:1000 Bittium 20808) CF647 (1:1000 Bittium 20808) secondary antibody incubation for 2 hours at room temperature. Lastly, washed cells were mounted on precleaned 1 mm thick microscopy slides with VectaShield DAPI mounting medium.

### e. Image Acquisition

Hardware configurations and image acquisition were managed using the Micro-Manager software [44]. First, we confirmed the proper functioning of the CSD-ISM setup, resulting in a resolution improvement with a ratio of 1.45. This closely aligns with the theoretically expected value of approximately 1.41 (√2) [supplementary section 2]. Next, the custom-designed linear Amici prism was integrated into our system, and consequently the DSP software parameters were adjusted for spectral image acquisition. It was crucial to ensure that the spacing between adjacent confocal spots, controlled by the “Laser Pulse Number (LPN)” knob, allowed for dispersion without overlap. Additionally, the number of frames for image acquisition had to be set in both the DSP software GUI and Micro-Manager software.

Reference images are essential for calibration adjustments during image processing and help optimize scanning parameters for image acquisition. Key parameters, including spinning disk rotation speed, laser pulse count, and the number of frames, are set based on the sample’s SNR and the number of color channels required for detection. In addition, by denoting the spinning disk period (single scan cycle) as TS, we established two constraints: TR = TS / LPN and TD = TR / N. The “Auto Rate” button automatically adjusted these parameters to meet the requirements. the parameters for each phase were:

- <u>Parameters for spectral calibration measurements:</u> disk spinning speed was set to 1800 RPM (equivalent to the pulse rate of 360 Hz), and the following control parameters: TC_ON = 50-200 (depending on the illumination), TC_OFF = 15, TL = 8 µs, TA = 0.0 µs, LPN = 2, N = 250.
- <u>Parameters for image analysis testing, spectral ISM validation and biological samples:</u> the disk spinning speed was set to 1500 RPM (equivalent to the pulse rate of 300 Hz), and the following control parameters: TC_ON = 100, TC_OFF = 15, TL = 12 µs, TA = 0.0 µs, LPN = 2, N = 1000. For some images, we down-sampled to 250 and 500 frames in order to shorten acquisition times.

All images featured in this study were subjected to adjustments in brightness and contrast using Fiji (ImageJ) [45]. Contrast enhancement was executed via Fiji’s ‘Enhance Contrast’ function, employing a saturation threshold of 0.35%. Furthermore, the minimum display brightness was calibrated to 10% of the maximum pixel intensity to optimize visual clarity.

### f. Image Processing

All image analysis and the computation of the spectral ISM image from the raw data stack were conducted utilizing Python [46]. For enhanced image resolution, the Richardson-Lucy (RL) deconvolution [37,38] method was applied with 50 iterations, by employing the Deconvolution Lab2, RL50 Algorithm plugin within Fiji [45]. [For schematic diagram of image processing pipeline see supplementary section 6].

#### i. Spectral calibration and simulations

To calibrate the system’s spectral response, a series of steps were undertaken. Experimental PSF images from a range of fluorophores were collected (namely Cy5.5, AF647, AF568, AF546, AF532, AF488, and AF405), and theoretical emission spectra were sourced from the Semrock Database. The calibration process involved fitting recorded intensity distributions to the theoretical center of mass wavelengths using a 4th-degree polynomial. This fit was employed to simulate PSFs for other fluorophores, accounting for optical components such as the dichroic mirror, emission filter, and camera’s spectral quantum efficiency. Using this calibration, we could simulate the spectral PSFs of additional fluorophores based on their theoretical emission spectra, which are readily available in online spectra-viewers [47–49] (supplementary section 2 and figure S2).

#### ii. localizing illumination spots’ positions

To register the illumination spots’ locations, for each experiment we acquired a single-color (AF647) reference stack with the same acquisition parameters as in the spectral acquisition of the samples. The illumination spots’ locations were registered with subpixel precision using the ‘trackpy’ Python package [50]. These locations were used both for ISM pixel reassignment and spectral ROI cropping as shown in figure 2.

#### iii. Spectral to Color Assignment

fluorophores present in the samples were selected, and their spectral PSFs were simulated. ROIs were then defined around each illumination spot, allowing for the cropping of spectral ROIs from the image stacks, thereby enhancing the accuracy of further analyses.

The spectral profiles for analysis and calibration were obtained from the Semrock’s SearchLight™ spectra viewer [47] as follows:

- <u>Image analysis validation</u> - The spectra profiles of the 4 µm fluorescent beads are: green-FocalCheck Double Green Ring (505/515), red - FocalCheck Red Ring (580/605), and far red - FocalCheck Dark Red Ring (633/660).
- <u>Spectral CSD-ISM characterization and validation</u> - The spectra profiles of the 170 nm fluorescent beads are: green-FluoSpheres Yellow-Green (505/515), orange - FluoSpheres Orange (580/605), and deep red - Alexa Fluor 633 (633/660).
- <u>Biological samples</u> - The following spectra profiles were used: nuclei - DAPI (359/461), green fluorescent protein GFP-eGFP (475/509), Ras GTPase-activating binding protein (G3BP) - CF568 (562/583) and synaptotagmin-1 - CF647 (652/667).

#### iv. Spectral to color-channel decomposition

To decompose the spectral signatures into multi-color channels, Ordinary Least Squares (OLS) regression was applied to the cropped ROIs. Each spectral signature within the cropped ROIs was decomposed into the simulated PSFs corresponding to the known sample labels through OLS regression. The output of the OLS analysis provided a set of three linear coefficients for each simulated PSF, indicating the composition of each PSF along with its two gradients based on their influence within the spectral ROI. To construct a color ROI, these triplet coefficients were linearly multiplied with the simulated PSF that most closely resembled a Gaussian profile (in this case, AF647). This process also incorporated gradient adjustments to account for minor parallax shifts, allowing for precise alignment. By utilizing the first Taylor approximation, the decomposition basis effectively enabled translation adjustments, thereby enhancing the accuracy of the reconstructed color image [see supplementary section 3].

#### v. Color and ISM image reconstruction

Following the decomposition of spectral signatures, the next step involved color image reconstruction. After each ROI was re-assigned both spatially and spectrally, the individual ROIs were summed and reconstructed to generate a multi-channel composite image of the sample. To enhance resolution, the ISM reconstruction was applied based on the algorithm developed by Qin [35]. Additional improvements were achieved through the Richardson-Lucy algorithm for deconvolution, implemented via a Fiji plugin. This comprehensive approach facilitated the generation of final full-color and ISM images, allowing for improved clarity and detail in the visualization of the sample.

## Notes

https://github.com/ebensteinLab

### Competing Interest Statement

The authors have declared no competing interest.

