## Supplementary Information for "A Spectral Image Scanning Microscope for Multi-Color High-Resolution Imaging"

1. **Prism Design:**

To optimize the color multiplexing capabilities of our system, we designed a direct vision prism (also called Amici prism) featuring linear dispersion across the visible spectrum. Previous studies utilizing refractive elements for spectral readout  [[25,36,51,52]](https://www.zotero.org/google-docs/?B8RdS7), typically employed standard prisms characterized by nonlinear dispersion, where the light dispersion depends nonlinearly on its wavelength. In applications when no spatial reference for the spectral PSF is available, this nonlinearity can improve color classification by exploiting small wavelength-dependent variations in the spectral PSF [[31,32,36]](https://www.zotero.org/google-docs/?IWtdgj). However, this comes at the cost of non-efficient color registration. As previously demonstrated, the optimal spectral dispersion is achieved by the minimal spectral resolution that can differentiate between the two spectrally closest dyes used in a given experiment [[36]](https://www.zotero.org/google-docs/?4eHsUr). Lower spectral resolution cannot adequately resolve colors, while excessively high resolution disperses spectral signatures over larger pixel areas, causing overlapping of adjacent spectral PSFs and thus reducing throughput. Furthermore, the extended pixel area caused by larger dispersions degrades the signal-to-noise ratio (SNR) due to the added pixels read noise. Nonlinear prisms inherently necessitate compromises: optimizing dispersion in one spectral region (e.g., blue) inevitably results in suboptimal dispersion in other regions (e.g., red). Consequently, the full spectral information density that could otherwise be encoded between adjacent excitation points is not fully utilized. In our system each excitation PSF is well defined spatially providing a spatial reference needed for unambiguous spectral readout. Therefore, linear dispersion ensures more efficient camera space utilization, enabling optimal spectral resolution per pixel and maximizing the density of encoded spectral information between adjacent excitation points.

To design a prism with linear dispersion, we utilized a custom optimization procedure implemented in Zemax. Due to the complexity of analytically solving Snell’s law in the absence of the paraxial approximation and with wavelength-dependent refractive indices, Zemax provided a powerful computational approach. We optimized the selection of prism materials and apex angles (n1, n2, α and β in figure S1-a respectively) by employing a merit function that balanced spectral resolution with dispersion linearity. To quantify and minimize the nonlinearity of the dispersion δ(λ), two terms introduced by Hagen and Tkaczyk [[53]](https://www.zotero.org/google-docs/?CpCq4w) were optimized as custom operands to our merit function: **(i)** the spectral sampling ratio (SSR, eq. 1), assessing the ratio between the dispersion gradient minimum and maximum, measuring the uniformity of dispersion across wavelengths; and **(ii)** the dispersion nonlinearity (NL, eq. 2), quantifying deviations from ideal linear dispersion by calculating the deviation of the deviation of dispersion gradient from a constant. By iteratively adjusting materials and apex angles based on these criteria, we successfully designed an Amici prism exhibiting linear dispersion tailored specifically for our spectral imaging needs (see figure S1-b and supporting Zemax file and merit function). The prism design was manufactured by Shanghai Optics (after a few iterations due to manufacturing and stock materials limitations) and integrated in our system at the output of the Yokogawa CSU.

1. $SSR= \frac{max(|\frac{d\delta}{d\lambda}|)}{min(|\frac{d\delta}{d\lambda}|)}$
2. $NL=\int|\frac{d^{2}\delta}{d\lambda^{2}}|d\lambda$

Using Zemax we simulated a minimal realization of the imaging system, placing the prism between a Nikon 60X planApo objective and a standard 200 mm tube lens (Figure S1-c). The Zemax files of the objective and tube lens were downloaded from online repositories and implemented in our simulation (objective from ref.  [[54]](https://www.zotero.org/google-docs/?6PcjrD) and tube lens as a black box file from Thorlabs’ website  [[55]](https://www.zotero.org/google-docs/?lor17u). The prism was optimized for: the availability of stock materials, total prism thickness, adequate separation between 5 color channels, and the camera’s pixel size. The optimization output was a linearly dispersing prism (Figure S1-d) which generated the optimal spectral PSF for separating 5 color channels while maintaining a minimal extension of the PSF (figure S1-e-f). The corresponding design was later manufactured by Shanghai Optics and is presented in figure S1-g.

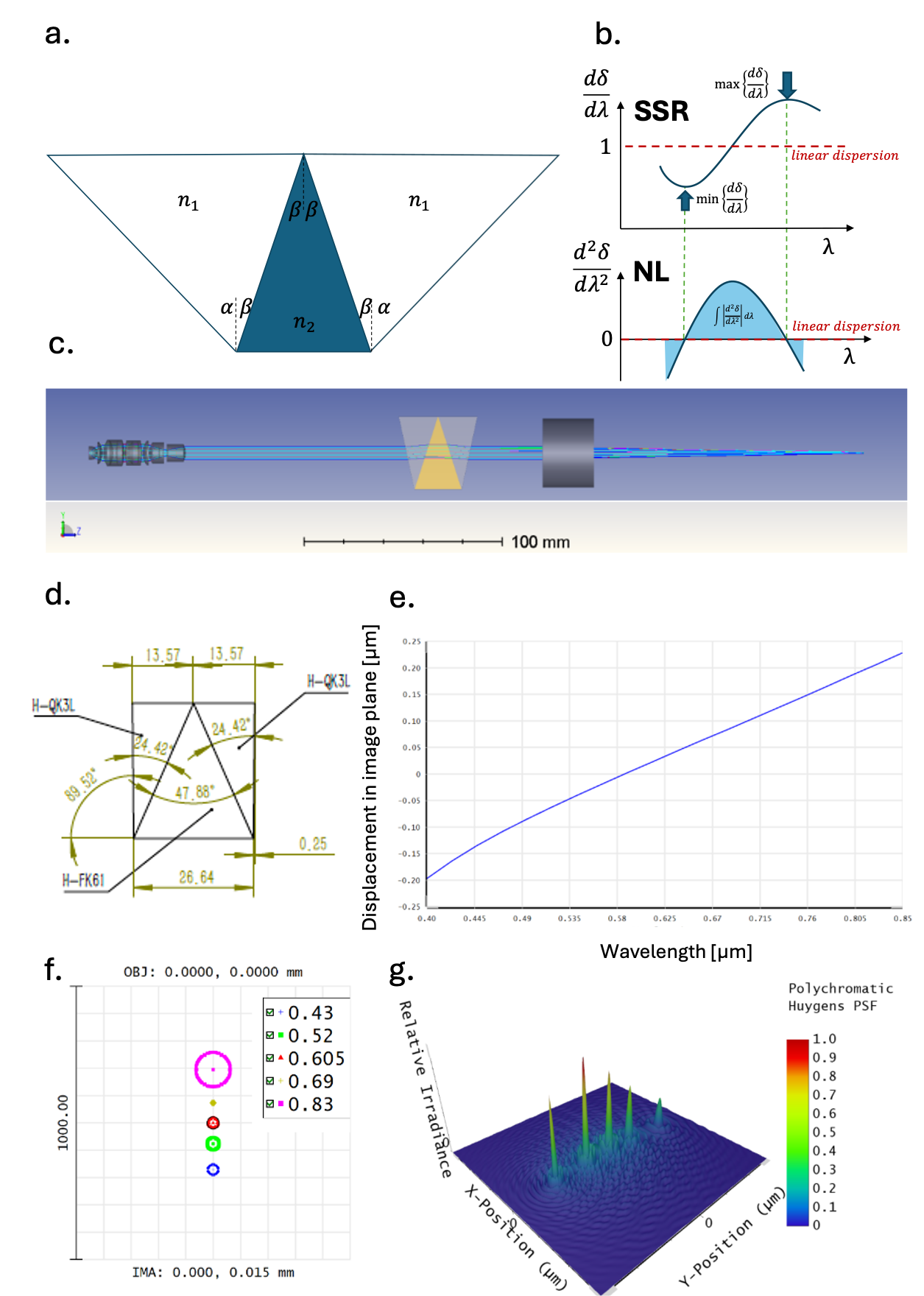

**Figure S1 – The design of a linear Amici prism**. **(a)** *Schematic illustration of the variables in the linear compound Amici prism optimization*. **(b)** *Graphical representation of the dispersion linearization criteria*. Linear dispersion is achieved when SSR=1, and NL=0. **(c)** *The layout of the Zemax optimization* using a 60X planApo objective and 200mm tube lens. We optimized the prism materials and apex angles for maximal displacement linearity in the image plane and minimal overlap of five spectral bands. **(d)** *The output design for an optimized Amici prism with linear dispersion*. **(e)** *Dispersion curve across the visible spectrum*, showing the linearity of the designed prism and its direct vision property (no beam deviation at 0.59 μm wavelength). **(f)** *A simulation of the spectral PSF* using the designed prism with five spectral bands. **(g)** *A physical ray simulation (Huygens) of the spectral PSF* showing the separation between the spectral bands in a realistic wave propagation.

2. **Spectral PSF Simulations**

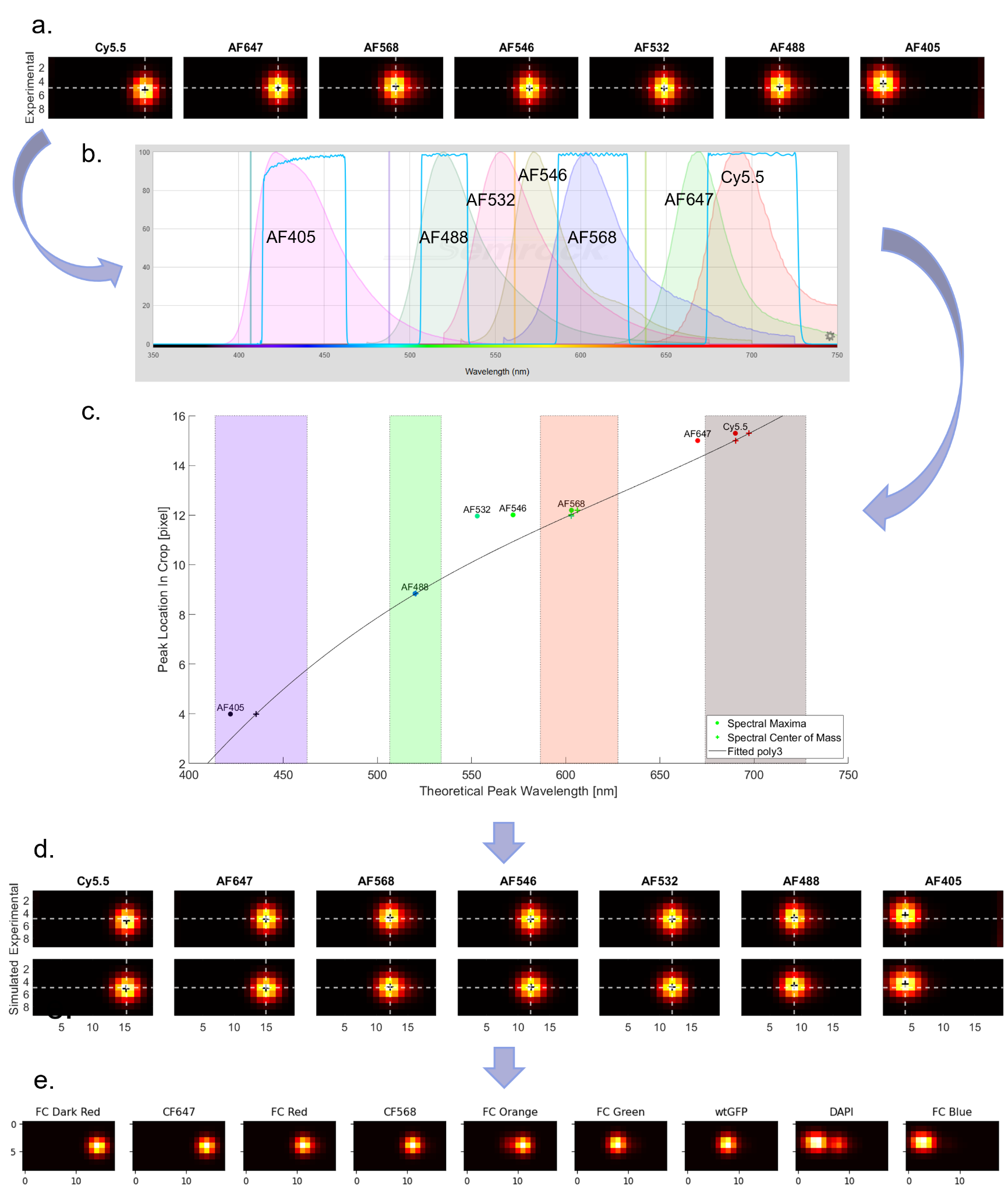

**Figure S2 – Pipeline for PSF Simulation.** Illustration of a comprehensive pipeline for simulating PSFs by integrating experimental data with theoretical models to achieve accurate PSF simulations for a variety of fluorophores. **(a)** *Acquisition of experimental PSF images from several fluorophores, namely Cy5.5, AF647, AF568, AF546, AF532, AF488, and AF405*. These images serve as the foundational data for subsequent analysis. **(b)** *Retrieving theoretical spectral data and filter information for each fluorophore from the Semrock Database.* The colored graphs depict the emission spectral profiles of the respective fluorophores, while the blue square graphs represent the transmission characteristics of our quad-band filter. The four vertical lines indicate the excitation wavelengths of the lasers used in the experiments. **(c)** *Plots and comparison of experimental PSFs to theoretical wavelengths*, employing a 4th-degree polynomial fit to map the wavelength to pixel locations. This calibration is essential for accurately simulating the system's spectral response. **(d)** Utilizing the polynomial fit, PSFs are simulated and validated against the experimental data, ensuring that the simulations are robust and reliable. **(e)** Simulate PSFs for unmeasured fluorophores, including FC Dark Red, CF647, FC Red, CF568, FC Orange, FC Green, wtGFP, DAPI, and FC Blue. This step is critical for expanding the applicability of the PSF simulations across a broader range of fluorophores.

3. **Spectral to color-channel decomposition**

The resolution enhancement achieved with ISM originates from the ability to image the sample from multiple viewing angles (parallax) and then correctly re-assign pixel intensities to their correct physical origin. Therefore, special care should be taken to capture in our color-assignment algorithm the small translation shifts induced by the different microlens array positions recorded at each frame. Figure S3 demonstrates the parallax effect with a single 170 nm PS-Speck 505/515 bead captured at multiple positions of the microlens array. These small deviations from the spectral PSF’s nominal lateral position (set by the dispersion) can amount to 2 pixels in both lateral dimensions. While these parallax-induced translations are essential for the resolution enhancement, they add complexity to the spectral-to-color decomposition algorithm. Naive implementation of our algorithm using only the spectral PSFs as the basis for the linear regression, omits all recorded parallax translations and therefore deteriorates the resolution. On the other hand, large parallax translations in the x-axis can be interpreted as spectral shifts, inducing misclassification of the colors in the spectral ROIs.

To solve this problem, we used the first Taylor approximation to capture the small parallax translations of the spectral PSFs. As shown in eq. 3, the first order Taylor expansion allows approximating a two-dimensional function (in our case the spectral PSF with a lateral parallax-induced shift), using a linear combination of the non-shifted spectral PSF and its two-dimensional gradients.

3. $F(x_{0}+\Delta x,y_{0}+\Delta y)\approx F(x_{0},y_{0} )+\Delta x\frac{\partial F}{\partial x}│_{(x_{0},y_{0} )}+\Delta y \frac{\partial F}{\partial y}│_{(x_{0},y_{0} )}$

With this Taylor approximation approach, we use eq. 4 to model the spectral ROIs (sROI) as a linear combination of three matrices for each of the fluorophores labeling the sample: the spectral PSFs (sPSF*_i_*) and its gradients $\frac{\partial(sPSF_{i})}{\partial x} and \frac{\partial(sPSF_{i})}{\partial y}$ (where *i* stands for a specific label):

4. $sROI=\sum_{i} C_{i}\cdot(sPSF_{i})+\Delta x_{i}\cdot\frac{\partial(sPSF_{i})}{\partial x}+\Delta y_{i}\cdot\frac{\partial(sPSF_{i})}{\partial y}$

The coefficients multiplying the gradients hold physical interpretation as the small parallax-induced lateral shifts, while the coefficients multiplying each sPSF stand for its intensity contribution in the sROI thus allowing determining the color composition. Using the OLS linear regression with these sPSF and gradients as orthogonal linear basis, allowed us to extract $C_{i}$, $\Delta x_{i}$, and $\Delta y_{i}$ for each dye the sample was labeled with.

Subsequently, as shown in eq. 5, to reconstruct a non-dispersed color-channel ROI we multiplied each label’s extracted coefficients with the same gaussian PSF ($gaussPSF$ in eq. 5).

5. $channel_{i}=C_{i}\cdot(gaussPSF)+\Delta x_{i}\cdot\frac{\partial(gaussPSF)}{\partial x}+\Delta y_{i}\cdot\frac{\partial(gaussPSF)}{\partial y}$

This resulted in a non-dispersed color-channel ROI reconstruction (channel*_i_* in eq.5) maintaining approximated parallax shifts required for resolution enhancement.

Figure S4 shows the results of the OLS color decomposition algorithm with parallax shifts registration through Taylor approximation. As can be seen in figures S4 and S5, the reconstruction of the spectral ROIs using this approximation is limited to small lateral shifts. When the shifts are large compared to the sPSF’s width they cannot be accounted for by the gradients alone.

Extending our approach to the second order of the Taylor approximation does not solve this problem. Introducing the second derivatives (figure S5) to the linear basis of our OLS algorithm results in failure of the color classification due to mixing the contributions of spectral dispersion and parallax-induced lateral shifts on the x-axis spectral ROI intensity distribution. Constraining the coefficients of the second derivatives in the regression could alleviate this issue, allowing to favor more physically plausible solutions. However, to the best of our knowledge, unlike the unconstrained OLS algorithm there is no closed-form calculation allowing to constrain the regression coefficient solution. Therefore, an iterative optimization is required to apply a constrained second order approximation, significantly adding to the computational cost and leads to significant performance deterioration when applied to many thousands of spectral ROIs. Comparing the computation time between the two approaches showed that the OLS was more than five orders of magnitude faster in decomposing the spectral ROIs, where the entire dataset was completed in a fraction of a second calculated on the CPU, whereas box-constrained algorithms took hours to complete decomposing the same dataset, although with slightly better reconstructions.

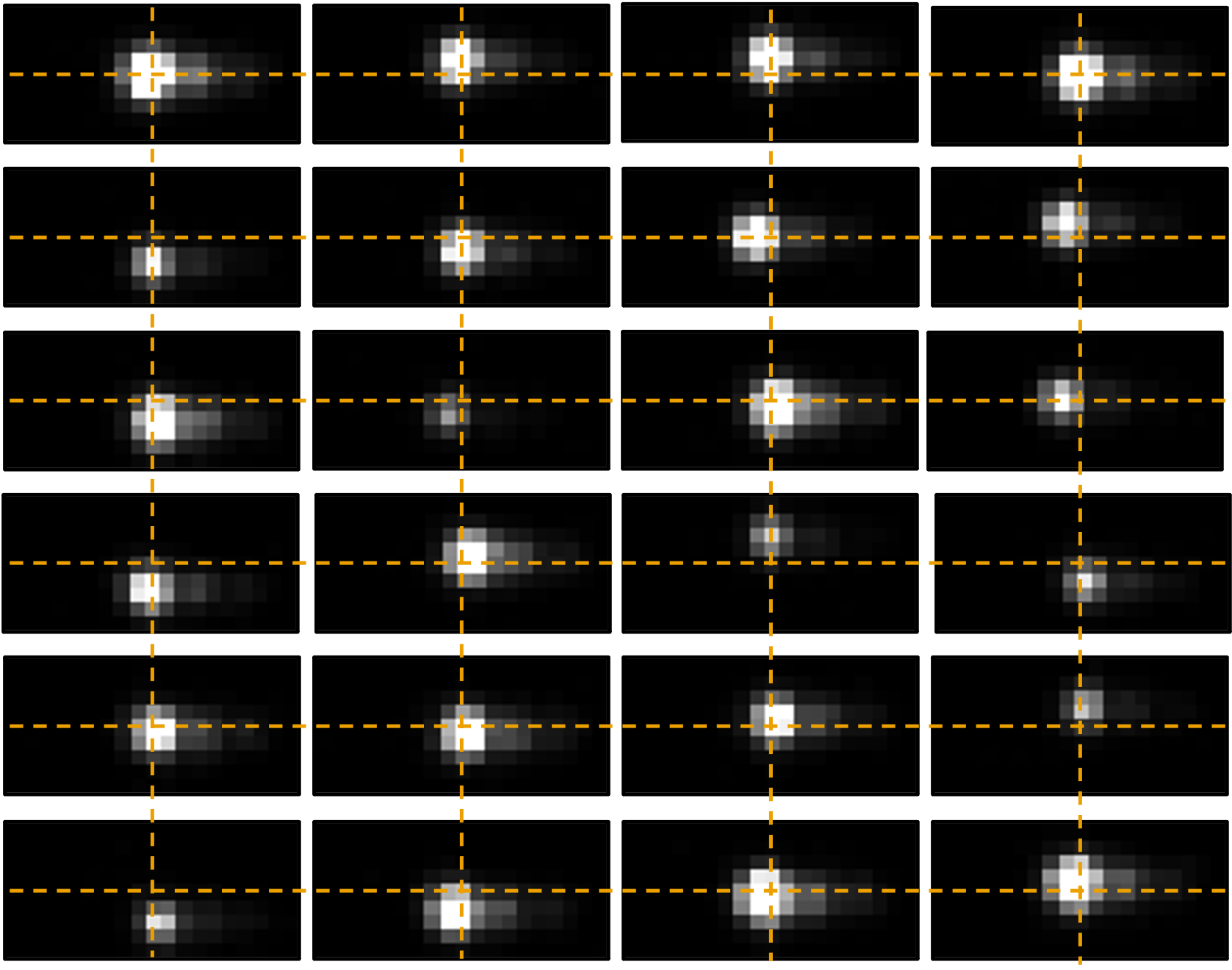

**Figure S3 – Small paraxial spatial shifts induced by imaging the sample through a variety of microlens positions and viewing angles.** The figure shows multiple ROIs of the same 170 nm PS-Speck 505/515 bead imaged at different frames and microlens positions.

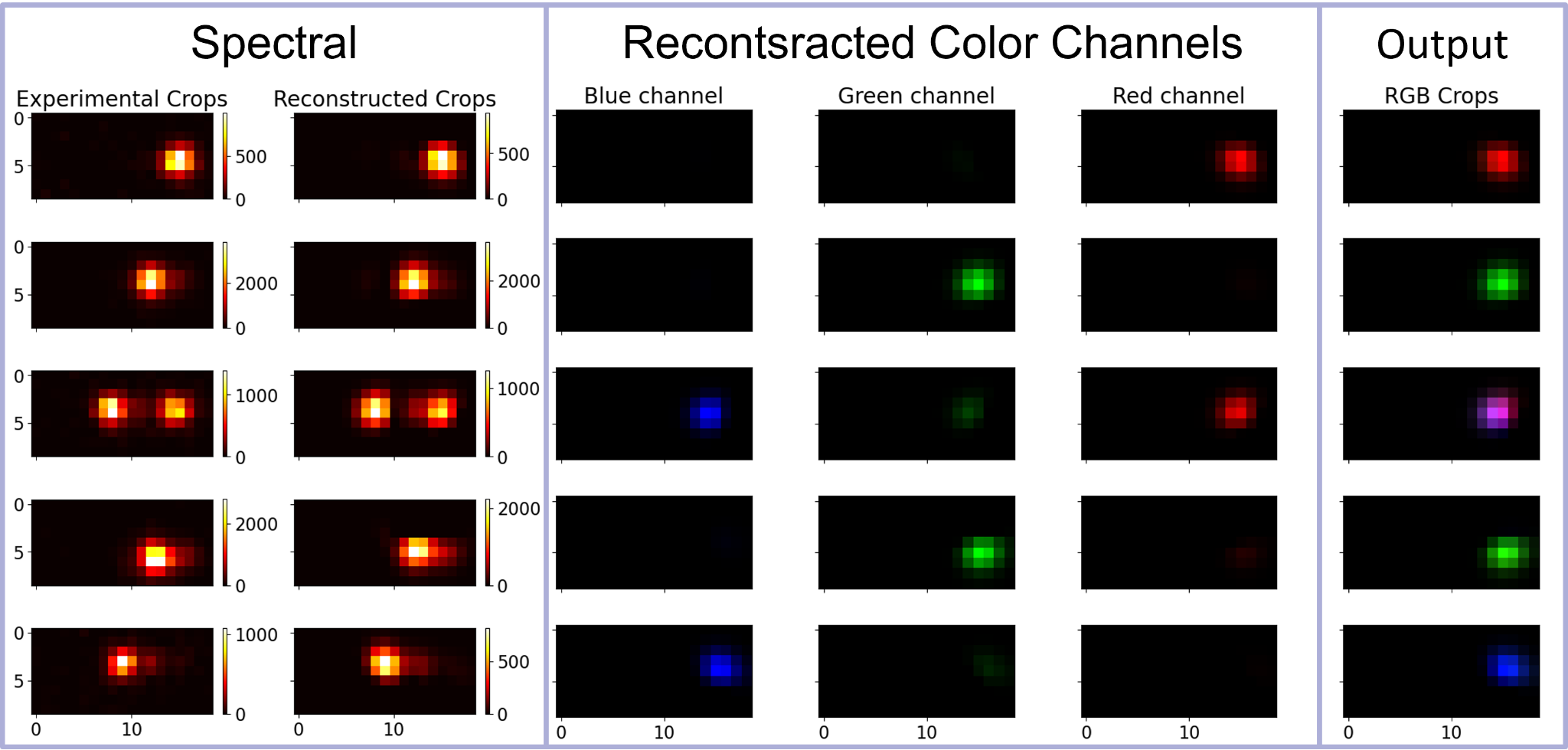

**Figure S4 – Spectral to color decomposition.** Spectral ROIs from a sample of three single color 170 nm fluorescent beads with emission maxima at 515 nm (blue colored), 560 nm (green colored) and 660 nm (red colored), side by side with their reconstructed spectral signature according to equation 4. The corresponding color-channel composition calculated with equation 5 is presented in the middle and a summed RGB ROI as the final output to the right.

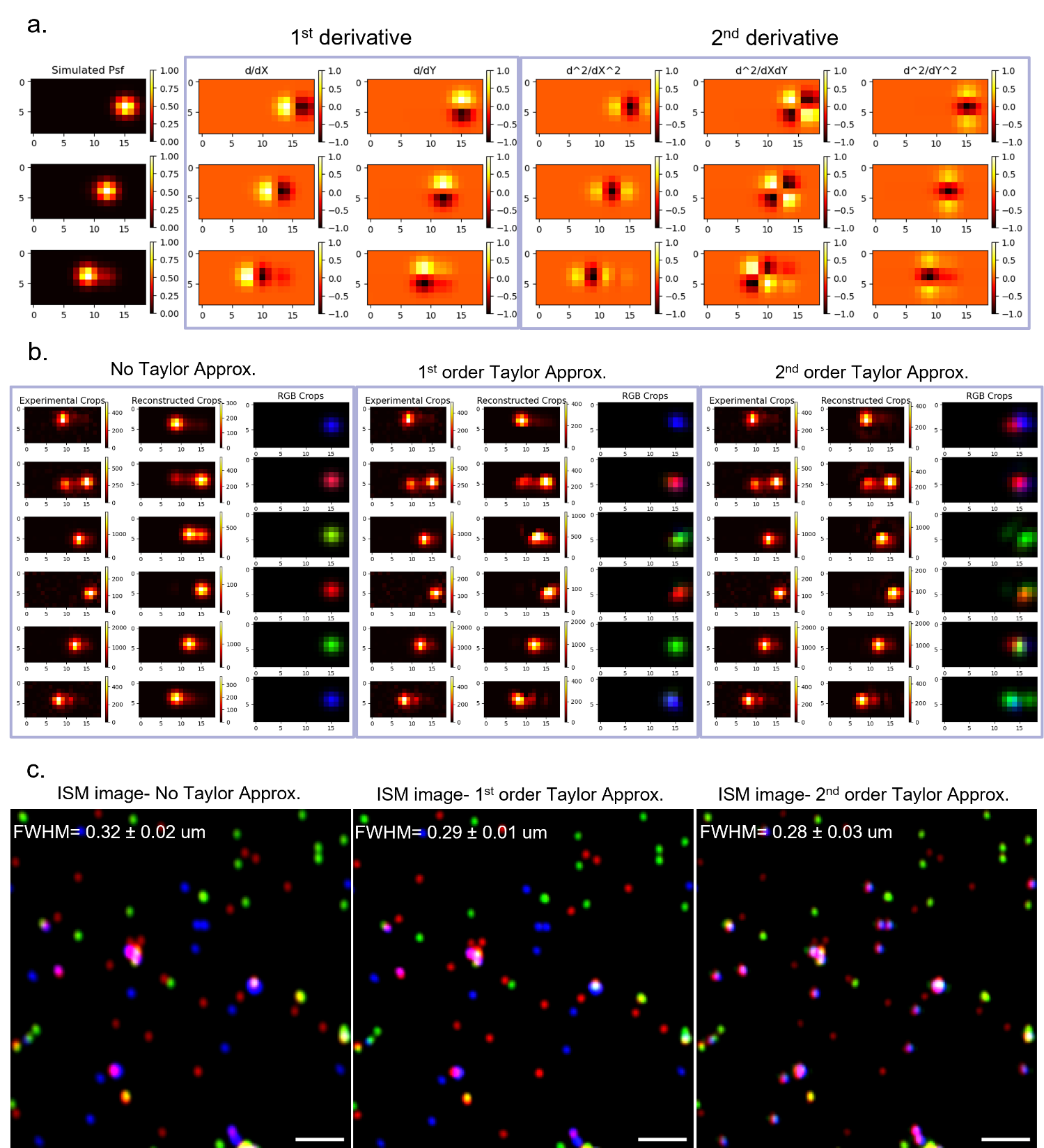

**Figure S5 – Comparison of image reconstruction and resolution using different Taylor approximation orders for parallax correction and color decomposition.** **(a)** *The sPSFs and their first and second-order spatial gradients used in the reconstruction process*. **(b)** *Comparison between reconstructions of individual fluorescent beads’ spectral ROIs*. The same beads’ spectral ROIs were reconstructed using different Taylor orders, for each order are shown from left to right: the spectral ROIs of the original experimental data, the reconstructed spectral ROIs calculated by the linear combination of basis images, and the reconstructed non-dispersed RGB color channels calculate with eq. 5. While the first order approximation manages to better reconstruct the parallax-induced translations, it is still limited in its reconstruction accuracy. Whereas the second order approximation introduces better spectral ROI reconstruction with deteriorated color classification. **(c)** *Images of fluorescent beads reconstructed using different methods or parameters*, with the measured FWHM values of the red beads indicated above each image as a quantitative measure of resolution. The decreasing FWHM values from left to right suggest improved resolution with the applied reconstruction techniques. However, the second order approximation introduces inaccuracies, causing the misclassification of color information. Scale bar, 2 μm.

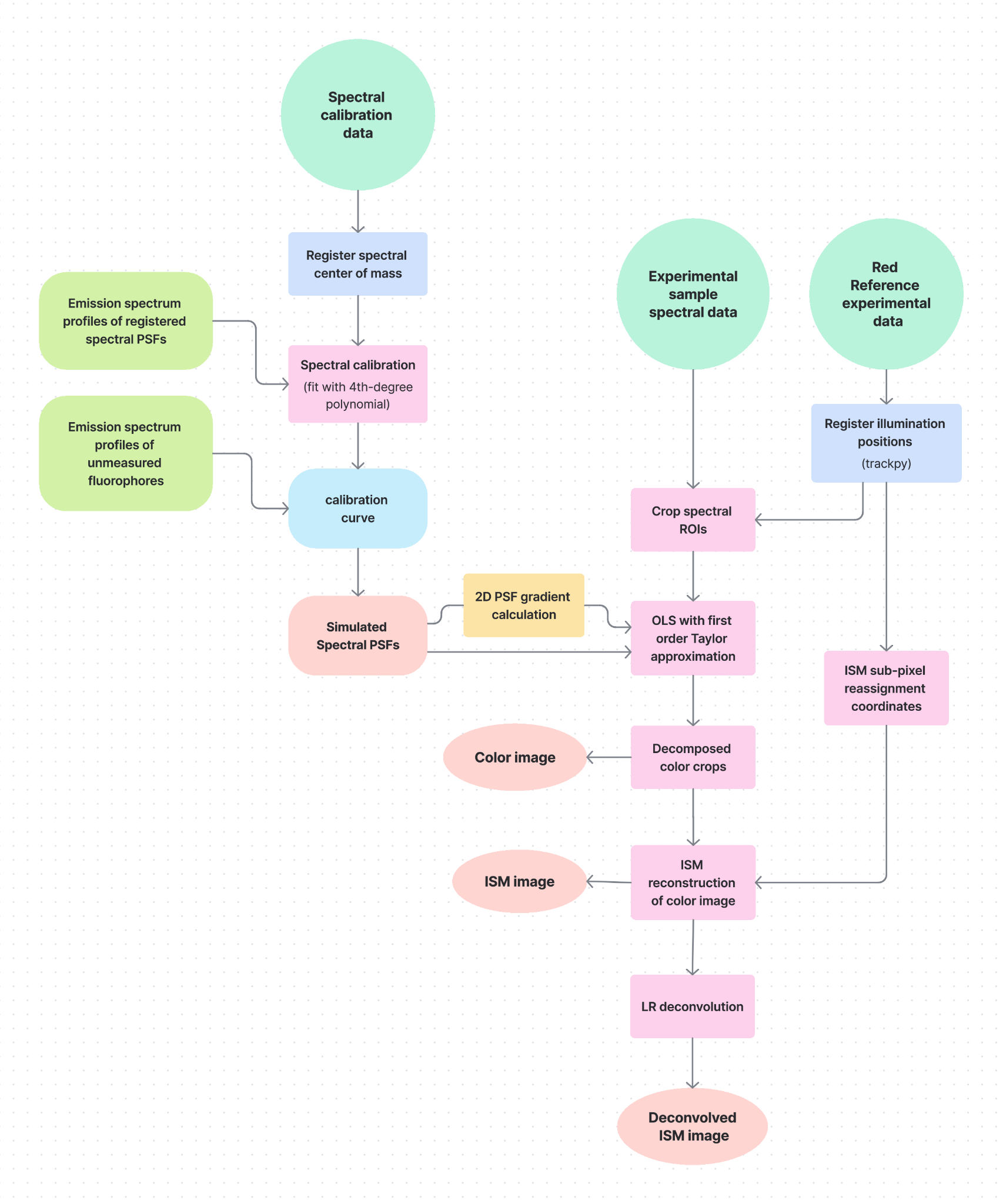

**Figure S6 –Flowchart of the analysis pipeline.**

4. **CSD-ISM Validation:**

To evaluate the effectiveness of the DSP-based controller and the resolution enhancement, we conducted a validation experiment along with a resolution assessment. For recording CSD-ISM images, the disk spinning speed was set to 1800 RPM (equivalent to a pulse rate of 360 Hz), with the following control parameters: TC_ON = 60, TC_OFF = 15, TL = 8 µs, TR = 463.33 µs, TD = 1.85 µs, TA = 0.0 µs, LPN = 6, and N = 250. For the reconstruction of ISM images, we utilized software developed by Qin et al. [[35]](https://www.zotero.org/google-docs/?DrbVoO). Calibration measurements were performed using a homogeneous fluorescent sample (a highlighter marker smeared on a coverslip and covered with a slide), and reference images were obtained by capturing CSD-ISM images. We then tested resolution enhancement using 0.1 μm TetraSpack fluorescent beads (ThermoFisher), which were excited with a 561 nm laser. Figure S7 illustrates a comparison between a spectral image obtained by averaging 250 recorded frames and the reconstructed ISM images. The inset zoomed-in images of two adjacent beads, along with a plot showing a line cross-section through the beads, demonstrate improved resolution and better separation between them. For a more accurate comparison, we measured the FWHM of the beads in both images. The ratio of the mean FWHM value in the averaged image to that in the ISM image indicates the resolution improvement. In our case, the experimentally obtained ratio of 1.467 ± 0.087 is close to the theoretically expected value of √2 ≈ 1.41 [3,10,35]. For further enhancement of resolution, we used R-L deconvolution and achieved resolution enhancement of experimentally 2.111 ± 0.093, aligning with the doubling the resolution in this imaging technique [[3,10,35]](https://www.zotero.org/google-docs/?27Oilu).

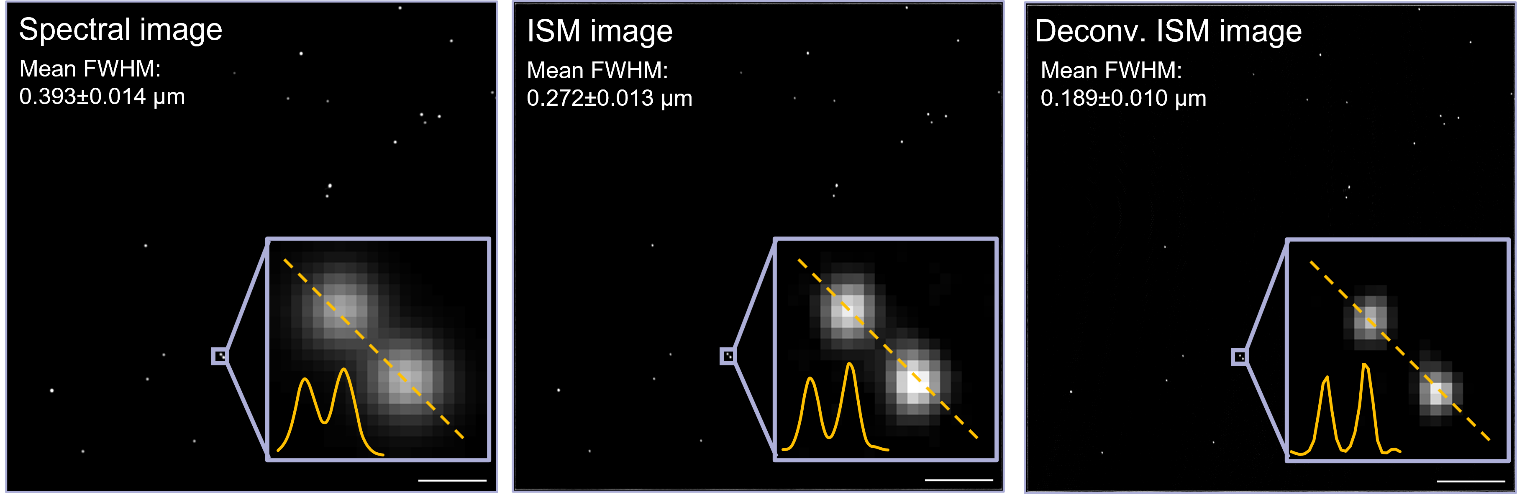

**Figure S7** – **Comparison of spectral averaged, ISM, and deconvolution ISM images of 100 nm fluorescent beads.** On the left is the spectral image, which has a mean FWHM of 0.393 ± 0.014 µm. The center shows the ISM image, demonstrating an improved mean FWHM of 0.272 ± 0.013 µm. On the right, the deconvolution ISM image achieves a mean FWHM of 0.189 ± 0.010 µm. scale bar; 10 µm. The zoomed-in insets across all three images illustrate the resolution enhancement, showcasing two neighboring and refined intensity profiles for each imaging technique, thereby confirming the significant improvement in resolution. The ratio of the mean values relative to the spectral image indicates a resolution enhancement of approximately 1.467 ± 0.087 for the ISM image and 2.111 ± 0.093 for the deconvolution ISM image, closely aligning with the theoretical expectations of √2 ≈ 1.41 and doubling of the resolution, respectively.

5. **Quantifying Resolution Enhancement:**

To quantitatively assess the resolution enhancement provided by different imaging modalities, including standard spectral imaging, color-coded representations, ISM, and deconvolution applied to ISM images, we employed 170 nm fluorescent beads as fiducial markers. Given the chromatic aberration observed in other channels with prism dispersion during standard spectral imaging, we focused our analysis exclusively on beads imaged at 633 nm (red channel). Particle locations were precisely identified via peak detection within the ISM image. For each detected bead, a one-dimensional intensity profile was extracted from a localized region centered on the particle in each of the four image types. The FWHM of the primary peak in these profiles was subsequently calculated, serving as a direct measure of the effective PSF width for each respective technique. By analyzing the distribution and average FWHM values across numerous beads for each modality, we quantitatively compared their resolution capabilities. Our measurements indicate a resolution improvement factor of 1.3 ± 0.16 for the ISM image and 1.74 ± 0.24 for the deconvolved ISM image, relative to the standard spectral imaging baseline. While previous studies  [[7,35]](https://www.zotero.org/google-docs/?Sr9MNZ) have reported resolution improvements of approximately √2 (≈ 1.41) for ISM and achieving nearly double the resolution of standard techniques after deconvolution, our observed improvement factors are somewhat lower. This discrepancy is attributed to the specific color decomposition and deconvolution algorithms employed in this study.

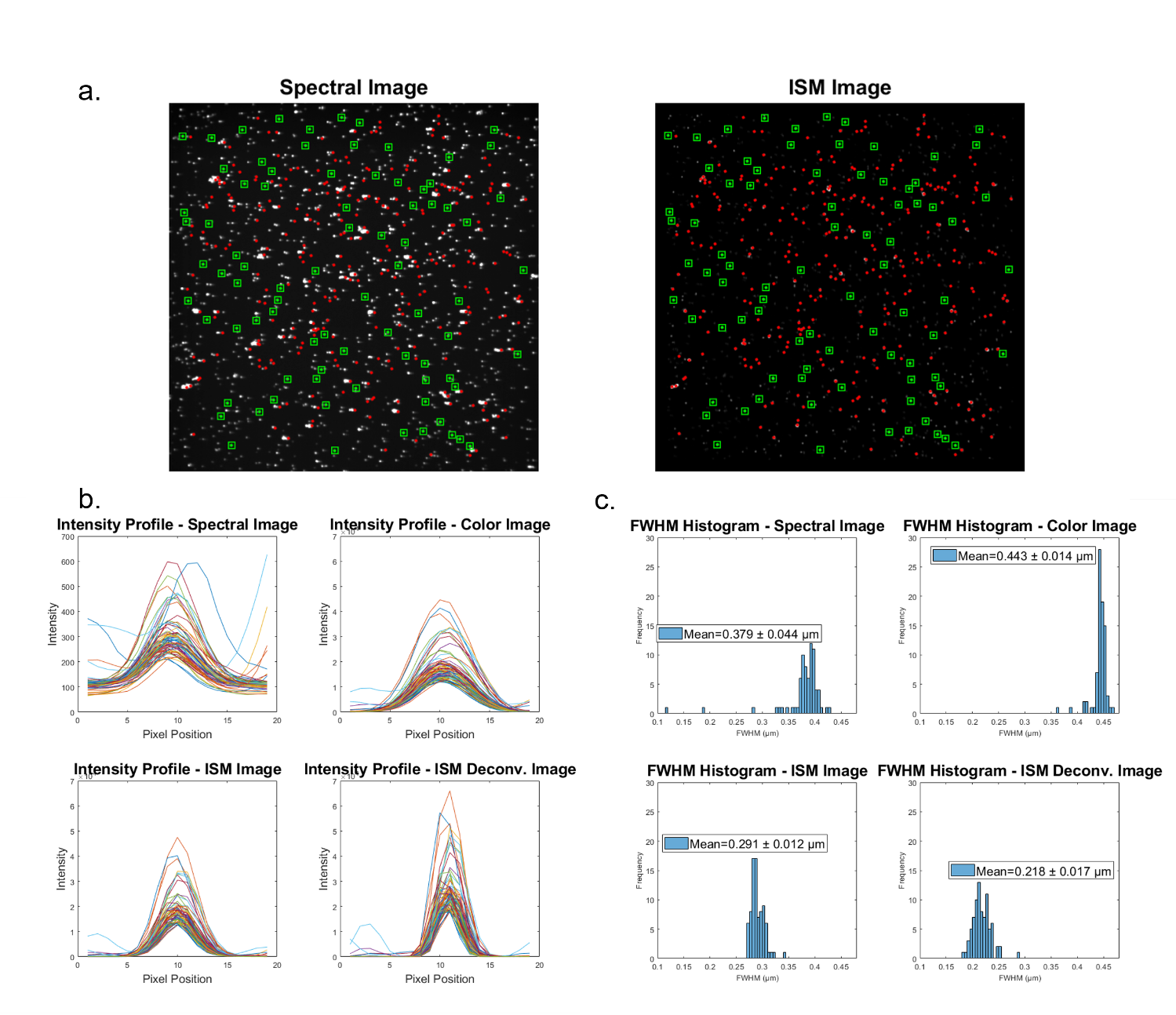

**Figure S8 – Quantitative Resolution Assessment of Different Imaging Modalities.** Quantitatively assessing the resolution of different imaging modalities using 170 nm fluorescent beads. **(a)** *Representative images used for analysis*: a Spectral image and an ISM image. Green squares indicate the locations of detected fluorescent beads, identified via peak detection (in red) primarily within the ISM image. **(b)** *One-dimensional intensity profiles* extracted from localized regions centered on detected beads in four different image types: spectral image, color-coded image, ISM image, and deconvolved ISM image. These profiles were used to calculate the Full FWHM. **(c)** Histograms of the calculated FWHM values across numerous analyzed beads for each of the four imaging modalities.

6. **Image comparison with different frame rates:**

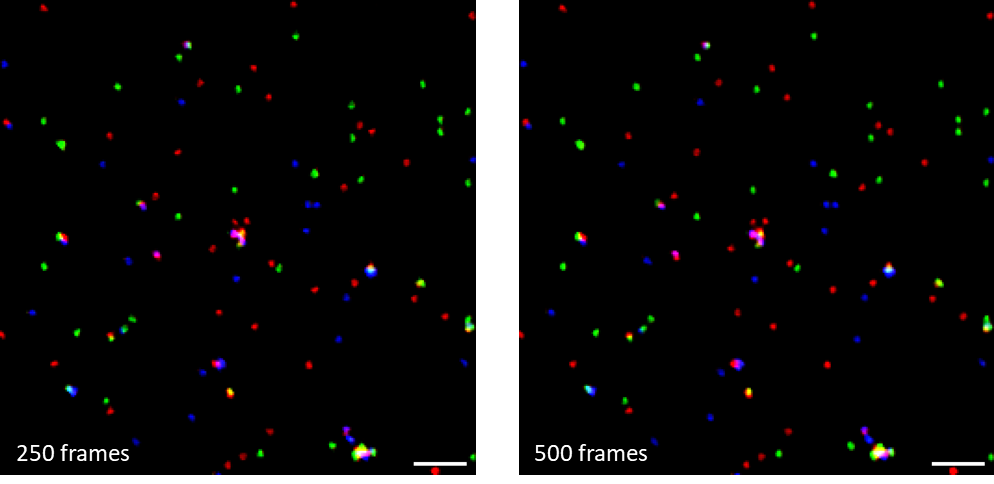
**Figure S9 – Image comparison with different frame rates demonstrating improved image quality with increased averaging.** Representative images of 170 nm fluorescent beads. The left panel shows a deconvolved single-color, non-dispersed, ISM image generated from a spectral image with a sum of 250 frames. The right panel shows the same region imaged with a sum of 500 frames. Note the deformation of the beads in the 250-frame image compared to the 500-frame image, highlighting the impact of frame averaging on image quality. Scale bar, 2 µm.

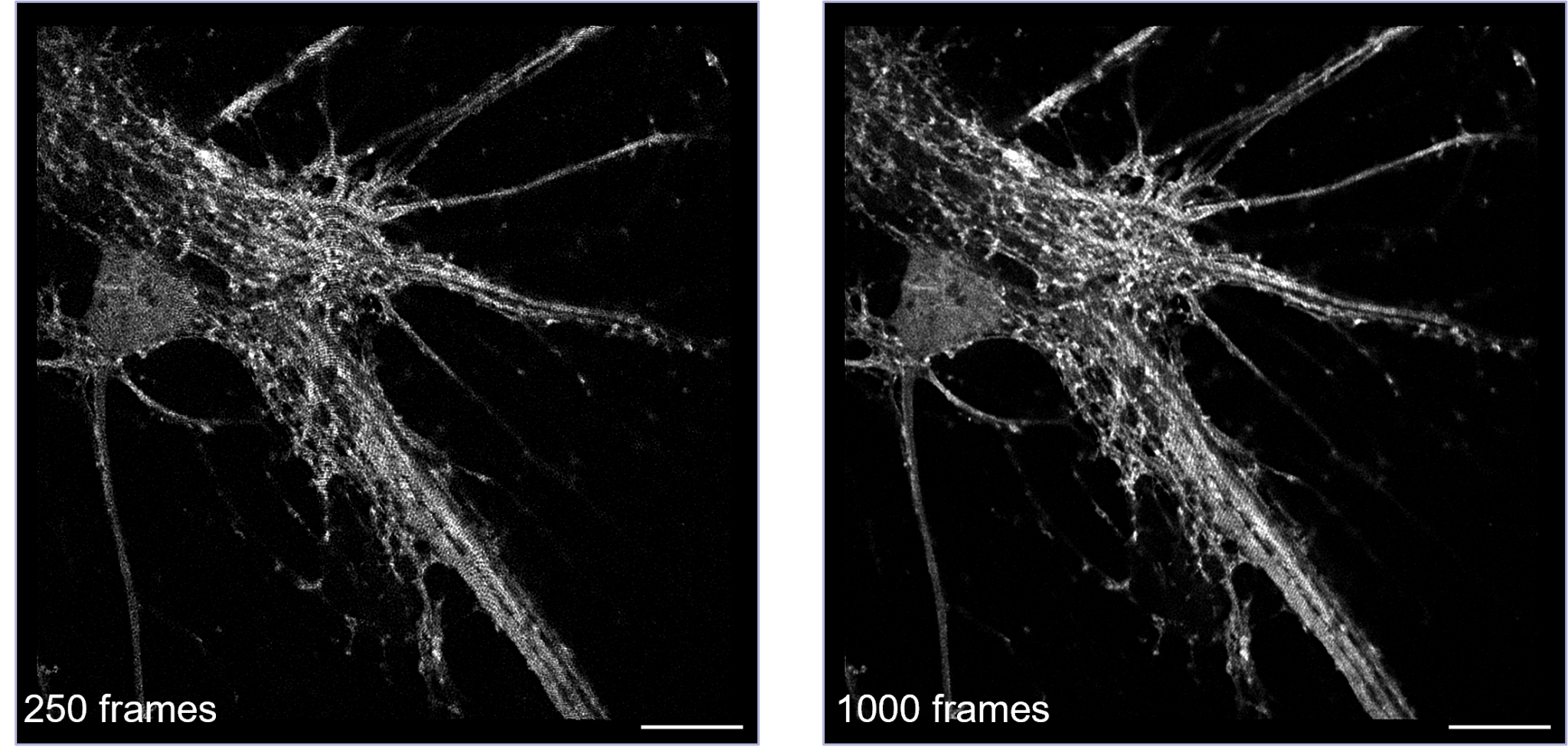
**Figure S10 – Image comparison with different frame rates, demonstrating improved image quality with increased averaging.** Representative images of adeno-associated virus endogenous eGFP expression in a neurite portion of a neuronal cell. The left panel shows a deconvolved single-color, non-dispersed, ISM image generated from a spectral image with a sum of 250 frames. The right panel shows the same region imaged with a sum of 1000 frames. Note the reduction in artifacts and improved SNR in the 1000-frame image compared to the 250-frame image, highlighting the impact of frame averaging on image quality. Scale bar, 10 µm.

7. **3D images:**

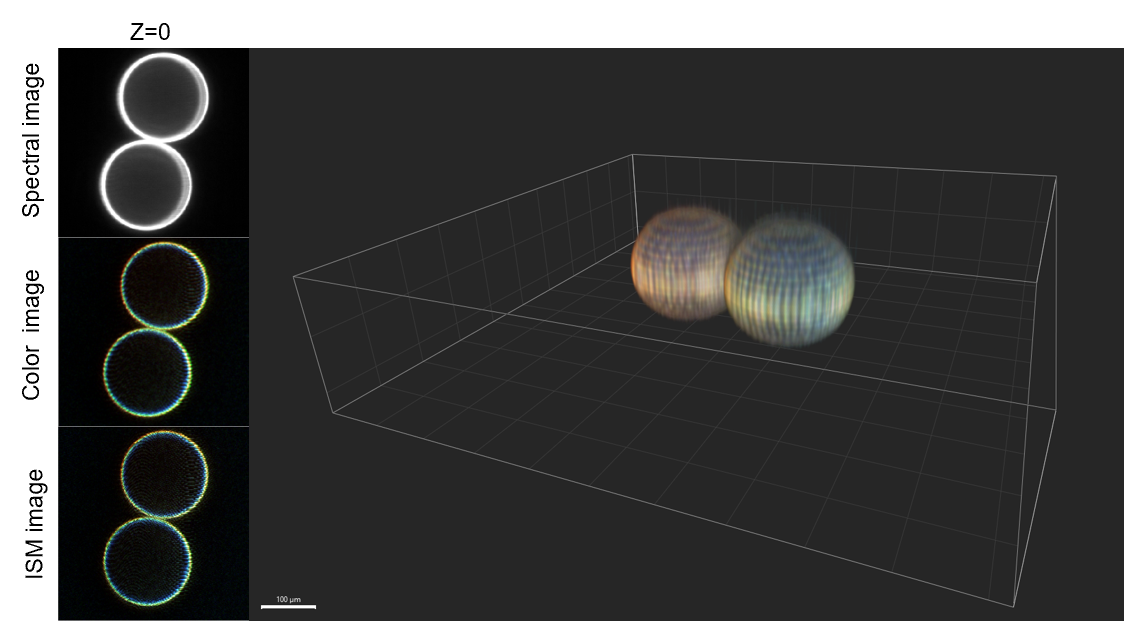

**Figure S11 – Fluorescently Labeled 15 µm Beads in 3D Visualization.** This figure illustrates a 3D representation of two 15 µm beads, each coated with three different fluorescent dyes, resulting in distinct thin rings of green (505 nm), orange (540 nm), and far-red (633 nm) fluorescence on their surfaces. The right side of the figure presents a 3D rendering of the beads, after spectral ISM image reconstruction, emphasizing their spherical shape and the spatial arrangement of the fluorescent coatings. On the left side, the images correspond to the Z=0 plane of the beads. The top image is the sum of 150 spectral images (spectral), middle, the sum of color assigned image without ISM reassignment (color) and the bottom, color assigned with ISM pixel reassignment (ISM).
